# Bigger is not always better: Optimizing leaf area index with narrow leaf shape in soybean

**DOI:** 10.1101/2025.07.07.663573

**Authors:** Bishal G. Tamang, Gregory Bernard, Carl J. Bernacchi, Brian W. Diers, Elizabeth A. Ainsworth

## Abstract

Modern soybean varieties have higher than optimal leaf area index (LAI), which could divert resources from reproductive growth. Altering leaf shape could be a simple strategy to reduce LAI. To test this, we developed 204 near-isogenic soybean lines differing in leaf morphology by introgressing narrow-leaf alleles from donor parents PI 612713A and PI 547745 into the elite, broad-leaf cultivar LD11-2170. We evaluated the lines across two locations and two row spacings (38-cm and 76-cm) to assess how reduced investment in leaf area influences canopy architecture, crop physiology, and yield. Narrow-leaf lines showed 13% lower peak LAI and 3% lower digital biomass compared to broad-leaf counterparts yet maintained yield parity (5,756 vs. 5.801 kg ha^-1,^ p = 0.43) across environmental conditions. Photosynthetic capacity remained largely unchanged, with narrow-leaf lines showing modest increases in electron transport rate and leaf mass per area. Narrow-leaf lines achieved similar canopy closure timing despite lower LAI, suggesting architectural compensation mechanisms. The most striking difference appeared in seed packaging, with narrow-leaf lines producing 34% four-seeded pods compared to only 1.8% in broad-leaf lines. There was a nonlinear relationship between peak LAI and yield, with optimal LAI values of 9-11 varying by environment. These findings show that the single-gene *GmJAG1*-controlled narrow-leaf trait offers a tractable strategy for reducing LAI and maintaining high productivity. This could reduce the metabolic costs associated with excessive canopy development and support sustainable agriculture under increasing climate variability.

## Introduction

Soybean (*Glycine max* (L.) Merr.) provides a critical source of protein, oil, and bioactive compounds for human consumption, animal feed, and industrial applications (Messina, 2022; soystats.com: accessed on 21 April 2025). Over the past few decades, global soybean production has expanded dramatically to meet increasing demand (FAO, 2023), but future production must address the dual challenges of climate change and limited arable land expansion (Lu et al., 2021; Goulart et al., 2023). This necessitates developing varieties with improved resource use efficiency to maintain and enhance productivity without increasing environmental footprints, a priority explicitly highlighted in the Soybean Genomic Research Community Strategic Plan (2024-2028), which underscores the need for innovative breeding strategies to develop resource-efficient, climate resilient cultivars (Stupar et al., 2024).

Agricultural sustainability increasingly depends on optimizing crop architecture for maximal productivity per unit of resource invested. The Green Revolution increased yields through architectural modifications, including reduced plant height and altered leaf angles to improve light interception and input responsiveness (Sakamoto and Matsuoka, 2004). Today’s “ideotype breeding” approach designs plants with ideal architectural and physiological traits for specific environments and cropping systems. Within this framework, leaf morphology in soybean represents a fundamental yet underexplored component of plant architecture with significant potential for optimizing resource allocation and enhancing yield potential (Roth et al., 2022, Clark and Ma, 2023, Li et al., 2024).

Plant architecture profoundly influences crop performance by determining spatial patterns of light interception, gas exchange and resource allocation (Nobel et al., 1993; Croce et al., 2024). In field conditions, individual plants function collectively as a canopy, where architectural traits dictate the efficiency of capturing available light energy for photosynthesis. For instance, the vertical distribution of leaf area and leaf angle controls light penetration, with upright leaves allowing greater illumination of lower canopy layers and enhancing whole-canopy photosynthesis (Sarlikioti et al., 2011; Mantilla-Perez and Fernandez, 2017; Yang et al., 2023, Sreekanta et al., 2024). Architectural traits also shape microclimates which affect water use efficiency, and they orchestrate nutrient utilization through distribution of photosynthetic capacity and nitrogen allocation within the canopy (Moreau et al., 2012). Further, modern crop modeling approaches increasingly incorporate detailed features of canopy architecture to capture these complex interactions. As climate variability increases, architectural traits that optimize physiological processes will become even more critical for maintaining crop production (Long et al., 2015; Liu et al., 2021).

Leaf Area Index (LAI), defined as the ratio of leaf area to ground area, serves as a critical indicator of canopy development and potential productivity (Parker, 2020; Wei et al., 2023). Optimal LAI varies by crop species, environment, and management, with both insufficient and excessive values of LAI potentially reducing yield. Insufficient LAI reduces light interception and photosynthetic capacity, while excessive LAI can lead to self-shading, reduced light use efficiency, and unnecessary allocation of resources to vegetative growth at the expense of reproductive development (Digrado and Ainsworth, 2023). For soybeans specifically, studies have identified nonlinear relationships between LAI and yield, with productivity peaking at optimal LAI values ranging from 3.5 – 6.5 depending on cultivar and growing conditions (Board and Harville, 1992; Liu et al., 2008; Setiyono et al., 2008; Tagliapietra et al., 2018). Research by Srinivasan et al. (2017) combining modeling and empirical approaches demonstrated that modern soybean varieties overproduce leaf area, a condition likely to worsen under future elevated atmospheric CO_2_ concentrations that increase biomass production (Ainsworth and Long, 2005; Soares et al., 2021). This nonlinear relationship suggests that maximizing leaf area may not be the optimal strategy for enhancing yield, particularly when considering the metabolic costs of leaf production and maintenance.

Plants face fundamental trade-offs in allocating limited resources between vegetative and reproductive structures. Leaves represent significant investment of carbon, nitrogen, and other nutrients, which must be balanced against investments in stems, roots, and reproductive organs (Holland et al., 2019). Leaf morphology – including size, shape, thickness, and arrangement – directly influences this resource allocation equation, affecting both the cost of leaf construction and the return on investment through photosynthetic output (Onoda et al., 2017, Villar et al., 2021). In crops, leaf shape diversity reflects both natural and breeding efforts. Breeding programs have traditionally focused on maximizing yield, which has indirectly increased leaf area. This approach may not optimize overall resource use efficiency. Narrow leaves potentially offer advantages in terms of reduced canopy structural investment, improved light penetration through the canopy and leaf-level photosynthetic rates (Tamang et al., 2023), which affects nitrogen distribution efficiency and photosynthetic nitrogen use efficiency (Qiang et al., 2023).

Recent advances in molecular genetics have identified key regulators of leaf morphology in soybean. The *GmJAG1* gene, encoding JAGGED-like transcription factor, has emerged as a critical determinant of leaflet shape. A single nucleotide polymorphism (SNP) within the EAR motif of *GmJAG1* causes an amino acid substitution (aspartic acid to histidine), disrupting its repressor function and resulting in a narrow leaf phenotype (Jeong et al., 2012). In Arabidopsis, the homologous *AtJAG1* promotes organ growth by repressing KIP-RELATED PROTEIN (KRP) genes, which inhibit the DNA synthesis phase of the cell cycle, thereby enhancing cell division (Schiessl et al., 2014). In soybean, *GmJAG1* likely operates similarly, with the non-functional allele reducing leaflet width by limiting cell proliferation, leading to elongated, narrower leaves. These morphological changes alter canopy architecture by reducing LAI and enhancing light penetration, potentially improving photosynthetic efficiency and resource allocation (Tamang et al., 2023).

Building on this genetic insight, our study leverages *GmJAG1’s* role to explore leaf shape’s broader impacts, addressing gaps in prior research. Despite some advances in understanding soybean leaf shape effects on canopy development and yield, existing studies have been limited in scale and genetic diversity. For instance, Bianchi et al. (2020) found narrow leaves improved light penetration but used only four isogenic lines, limiting statistical power and generalizability. Similarly, earlier work by Mandl and Buss (1981) and Wells et al. (1993) faced comparable limitations in genetic material. More recently, Cai et al. (2021) demonstrated yield benefits of narrow leaf soybeans through a gene editing approach, yet overlooked canopy dynamics. Our study overcomes these constraints by investigating over 200 near-isogenic lines that differ primarily in leaf shape, providing statistical power and genetic diversity to examine the relationships between leaf shape, LAI and productivity. Understanding these relationships provides valuable insights for breeding programs seeking to enhance resource use efficiency and yield stability in soybean in future climatic conditions.

The objectives of this study were to: 1. Characterize the effects of leaf shape on canopy development dynamics, including leaf area index, canopy coverage and biomass accumulation, 2. Assess whether altered leaf morphology affects photosynthetic capacity at leaf level, 3. Determine the impact of leaf shape on yield components and seed quality parameters, 4. Investigate how environmental conditions and management practices interact with leaf shape to influence plant performance, and 5. Identify key traits and processes mediating the relationship between leaf morphology and yield. To achieve these objectives, we developed and evaluated 204 near-isogenic soybean lines with a wide range of leaf shape across two environments and two management practices. Our phenotyping approach integrated detailed leaf-level measurements with canopy-scale assessments and yield evaluations, providing a holistic understanding of how leaf shape influences the source-sink continuum from photosynthetic capacity to yield.

## Results

### Isogenic lines differ in leaf shape, not phenology

Using marker-assisted repeated backcrossing, we developed 204 isogenic lines that significantly differed in leaf shape and LAI, but not most developmental or physiological traits (Figure 1A, B). Genotyping with 1327 Agriplex SNPs (829 polymorphic between parents) confirmed that the isogenic lines shared between 92 and 99% genetic identity with the broad-leaf recurrent parent LD11-2170 (Figure 2D), close to the theoretical similarity of 93.95% expected after three backcrosses. A *GmJAG1*-specific KASP marker effectively discriminated between homozygous broad-leaf, heterozygous, and homozygous narrow-leaf lines (Figure S4), while sequencing of the *GmJAG1* gene confirmed the presence of the causal SNP variant in the EAR motif among parental lines (Figure 2A).

**Figure 1.**
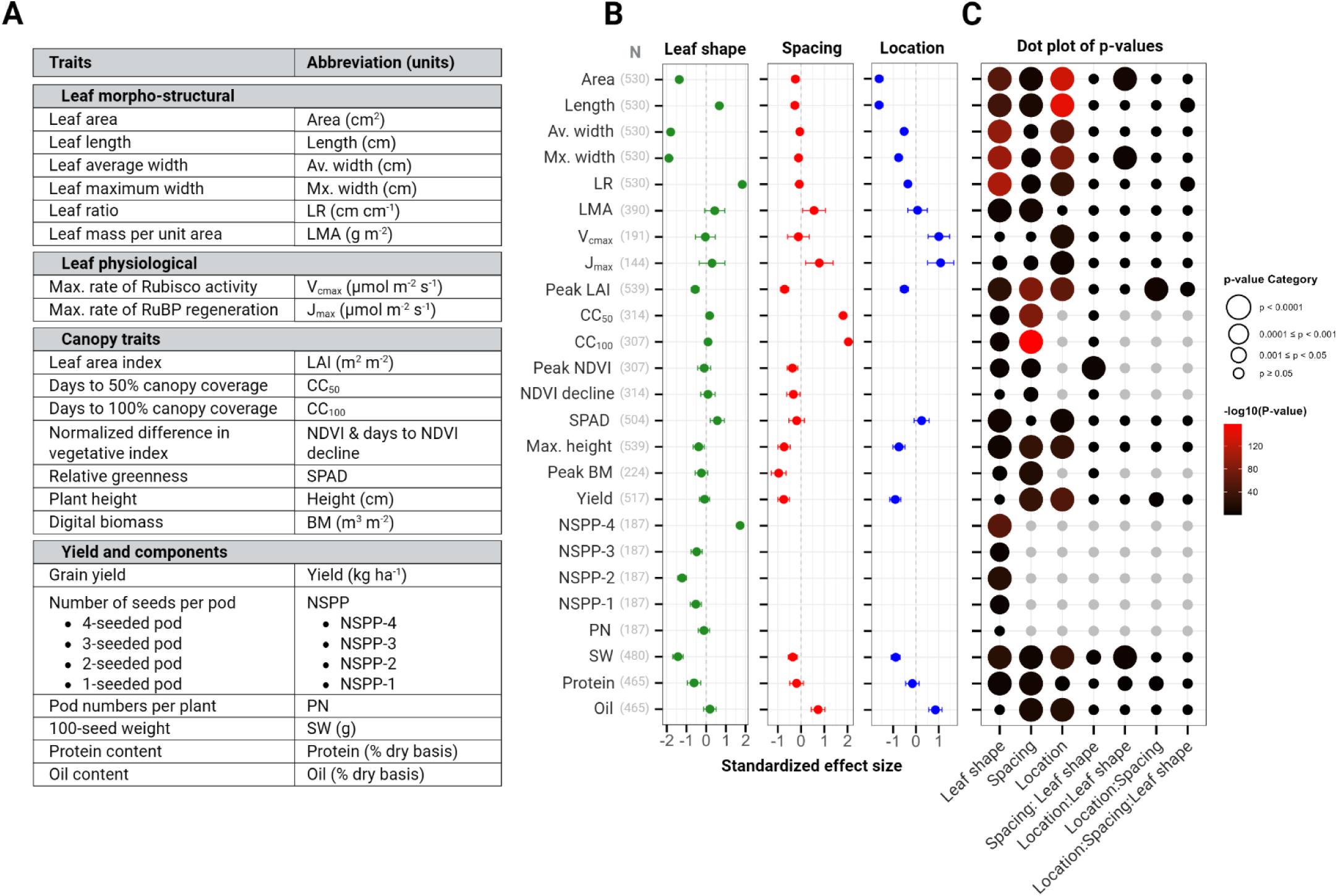
Mixed modeling results for trait responses to experimental factors. **A)** List of measured traits with their corresponding abbreviations and units of measurement. **B)** Standardized effect size (± 95% confidence intervals) from mixed models for each trait in response to Leaf shape (green circles representing effect of narrow leaf relative to broad leaf shape as reference, left panel), Spacing (red circles representing effect of 76-cm spacing relative to 38-cm spacing as reference, middle panel), and Location (blue circles representing effect of location SoyFACE relative to Energy Farm as reference, right panel). The y-axis displays trait abbreviations with sample sizes (N) used in the mixed models for each trait. **C)** Significance of main effects and interactions from mixed models. The x-axis displays the three main experimental factors (Leaf shape, Spacing and Location) and their four interaction terms (Leaf shape x Spacing, Leaf shape x Location, Spacing x Location, and Leaf shape x Spacing x Location). Circle size represents significance categories of p-values, while circle color intensity corresponds to −log_10_(p-value), with darker colors indicating higher statistical significance. All effects are relative to the respective reference levels (Broad leaf shape, 38-cm spacing and location Energy Farm). Positive effect sizes indicate that narrow leaves, 76-cm spacing, or SoyFACE location increased the trait value relative to their respective reference levels (broad leaves, 38-cm spacing, or Energy Farm location), while negative effect sizes indicate a decrease.

**Figure 2.**
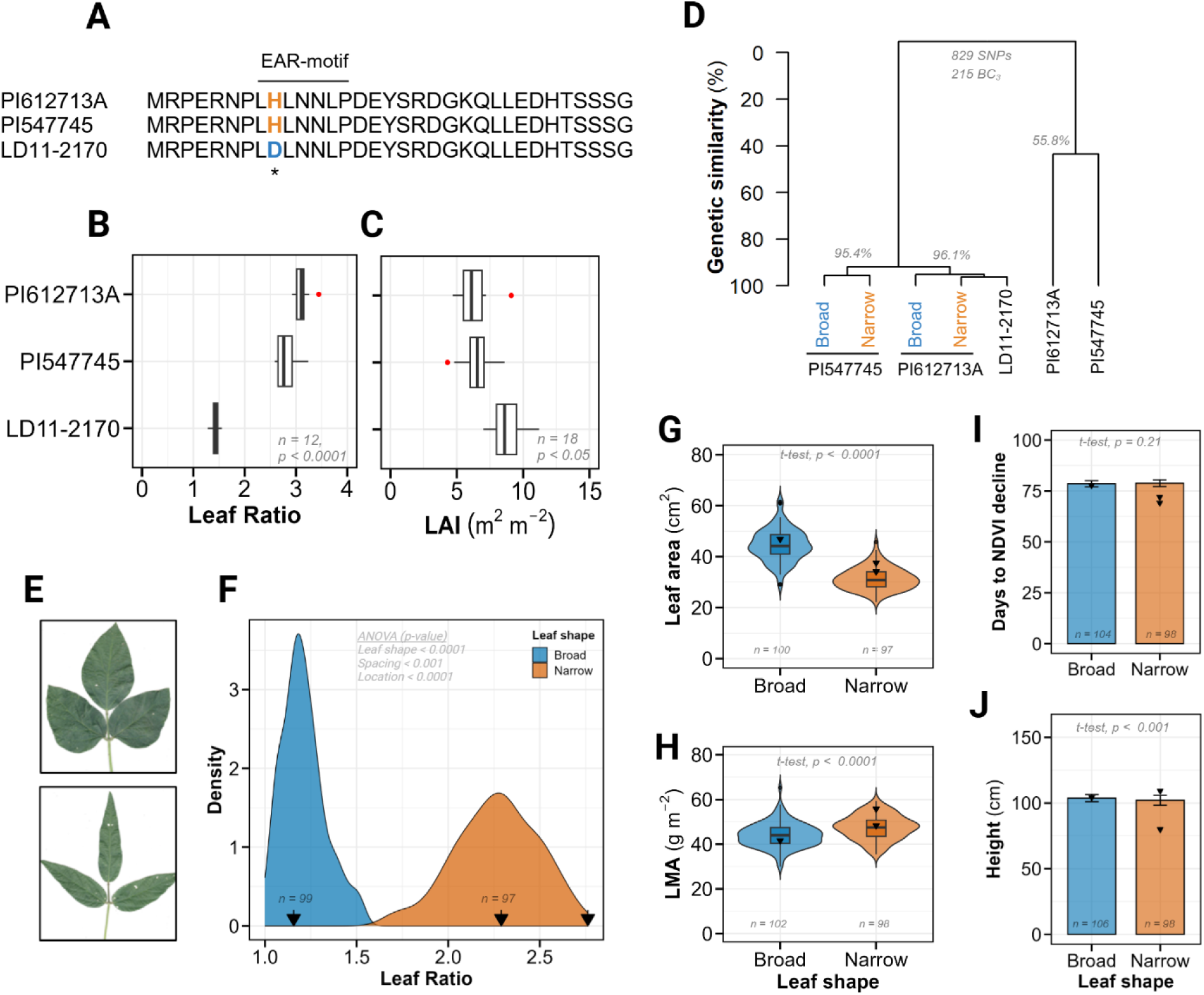
Molecular characterization and phenotypic differences of parental and isogenic lines. **A**) Amino acid sequence alignment of GmJAG1 from three parental lines: recurrent parent LD11-2170 and narrow leaf donor parents PI 612713A and PI 547745. The sequence displays the first 33 amino acids from the start codon (M), highlighting the critical single nucleotide polymorphism that causes an amino acid substitution from aspartic acid (D, blue) in LD11-2170 to histidine (H, orange) in both narrow leaf parents within the EAR motif region (indicated above the sequence) rendering the gene non-functional. **B-C)** Box plots comparing leaf ratio (B) and leaf area index (LAI, C) among the three parental lines. Plots display means, interquartile ranges, and minimum-maximum values, with red circles indicating outliers. Sample sizes (n) and ANOVA p-values are provided from each trait. **D)** Hierarchical clustering showing genetic similarity (%, y-axis) among the three parental lines and 204 BC_3_ progeny based on 829 polymorphic SNP markers. The BC_3_ progenies were classified into broad and narrow leaf categories derived from PI 547745 and PI 612713A, with percentage similarity values indicated at major branch points. **E)** Representative images of soybean trifoliate leaves from broad leaf (upper panel) and narrow leaf (lower panel) isogenic lines. **F)** Density plot of leaf ratio distribution for broad (blue) and narrow (orange) leaf phenotypes. Black arrows along the x-axis indicate the three parental values. Sample sizes (n) for each leaf shape category and significant mixed model ANOVA p-values are provided within the panel. **G-H)** Violin plots with embedded box and whisker plots comparing leaf area (G) and leaf mass per area (LMA, H) between broad and narrow leaf categories, Black triangles represent parental values, with sample sizes (n) and t-test p-values indicated. **I-J)** Bar plots showing days to NDVI decline (I) and plant height (J) between broad and narrow leaf categories. Black triangles indicate parental values, with sample sizes (n) and t-test p-values provided for each comparison.

The donor narrow-leaf parents (PI612713A and PI547745) exhibited distinct leaf morphology compared to the recurrent parent, with approximately two-fold higher leaf ratios (LR: 3.1 and 2.8 vs. 1.4, respectively) and substantially lower leaf area index (LAI: 6.3 and 6.5 vs. 8.8, p < 0.05; Figure 2B, C). The parents also differed significantly in LMA, height, V_cmax_ and J_max_ (p < 0.001, Figure S2B-E).

As intended, the resulting BC_3_ isogenic lines displayed a clear bimodal distribution in leaf ratio, with means of 1.3 and 2.4 for broad and narrow categories, respectively (Figure 2F). This leaf shape variation showed a large effect size (1.3, p < 0.0001; Figure 1B, C), indicating strong genetic control of this trait. The narrow-leaf phenotype was also associated with approximately 35% less leaf area (p < 0.0001) and 5% higher LMA (p < 0.0001) compared to broad-leaf lines (Figure 2G, H). Location also significantly influenced leaf ratio (effect size = −0.35, p < 0.0001), while row spacing did not (p = 0.16).

Despite these morphological differences, both leaf types showed similar developmental progression. Days to NDVI decline (a proxy for reproductive maturity) averaged 79 days for both leaf types (p = 0.21, Figure 2I), showing only marginal effects of spacing (effect size = 0.33, p = 0.026, Figure 1B-C). Similarly, plant height differences between leaf types were statistically significant but minimal in magnitude (108.3 cm for broad vs. 106.3 cm for narrow, p < 0.01; Figure 2J). These results confirm that our genetic intervention successfully altered leaf shape while maintaining comparable growth and development patterns, providing an ideal system to isolate the effects of leaf shape on canopy development and yield.

### Photosynthetic capacity is primarily determined by location, with minor influences from leaf morphology and row spacing

To investigate whether altered leaf morphology affected photosynthetic capacity, we analyzed CO_2_ response curves from 264 leaf samples representing all treatment combinations. Both maximum carboxylation rate (V_cmax_) and maximum electron transport rate (J_max_) varied strongly between locations, but did not vary as much by leaf shape or row spacing (Figure 1B-C). This suggested that photosynthetic biochemistry was not significantly altered by the changes in leaf morphology.

Location was the dominant factor affecting photosynthetic parameters, with large and significant effect size (1.0-1.1, p < 0.0001; Figure 1B). Both V_cmax_ and J_max_ were substantially higher at SoyFACE compared to the Energy Farm (approximately 17% and 12% higher, respectively; Figure 3A-D), likely reflecting differences in soil properties and/or microclimate between the two locations. These location-based differences were consistent across both leaf types and row spacings, underscoring the strong influence of environmental factors on photosynthetic capacity.

**Figure 3.**
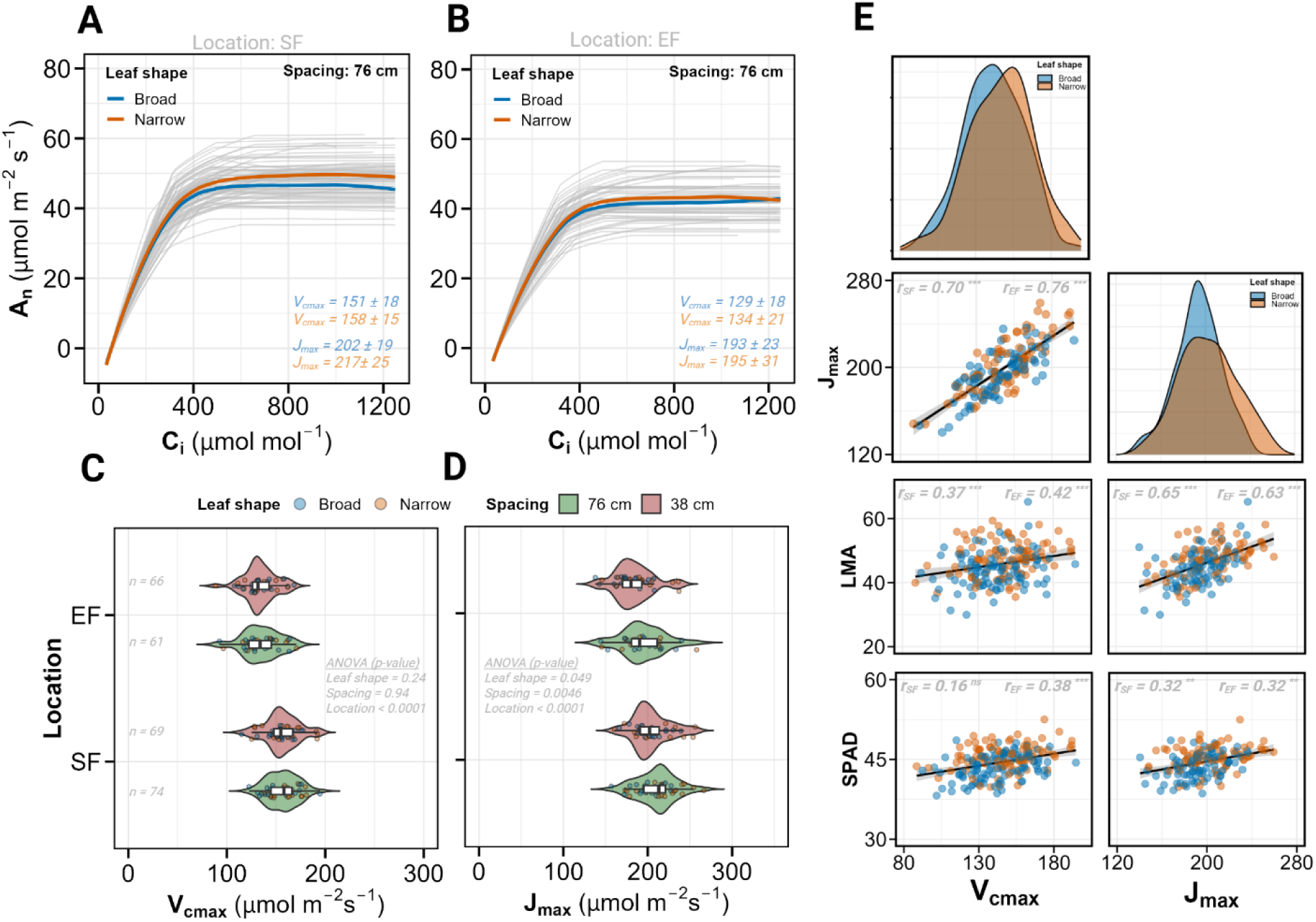
Photosynthetic parameters derived from A-Ci curve analysis and their correlation with leaf structure. **A-B)** Fitted A-Ci response curves from PhotoGEA R package for plants grown in 76-cm spacing at location SoyFACE (SF) (A) and Energy Farm (EF) (B). Curves represent model outputs from the package, which were generated at 1 µmol mol^-1^ C_i_ intervals based on fitted parameters derived from measured data points. Individual fitted curves are shown in light gray, with average curves for broad (blue) and narrow (orange) leaf shapes highlighted. Mean values (± SD) for maximum carboxylation rate (V_cmax_) and maximum electron transport rate (J_max_) are displayed for each leaf shape with corresponding colors. **C-D)** Violin plots with embedded box and whisker plots showing V_cmax_ (C) and J_max_ (D) distribution across locations (SF and EF) and row spacings (76-cm in green, 38-cm in red). Individual data points are colored by leaf shape (broad in blue, narrow in orange). Sample size (n) and ANOVA p-values from mixed models are provided. **E)** Pairwise correlation analysis between photosynthetic parameters (V_cmax_, column 1; J_max_, column 2) and leaf traits measured from the same leaf. The top panel shows density distributions by leaf shape for the two photosynthetic parameters. Remaining panels display scatter plots with regression line (black) and 95% confidence intervals (shaded areas) for relationships between the photosynthetic parameters and leaf mass per area (LMA), and SPAD. Data points are colored by leaf shape (broad in blue, narrow in orange). Pearson correlation coefficients (r) are provided for each location (SoyFACE and Energy Farm) with significance indicated by asterisks (* p < 0.05, ** p < 0.01, *** p < 0.001).

In contrast to the location effect on photosynthetic capacity, leaf shape had no significant influence on V_cmax_ (p > 0.2) and a small yet significant effect on J_max_ (effect size = 0.29, p < 0.05; Figure 1B, 3A-D). Narrow-leaf isogenic lines had ∼4% higher J_max_ compared to their broad-leaf counterparts, suggesting a slight enhancement in electron transport capacity. Similarly, row spacing showed no significant effect on V_cmax_ but modestly influenced J_max_ (effect size = 0.78, p < 0.05), with 76-cm spacing associated with 6% higher J_max_ compared to 38-cm spacing.

Correlation analysis revealed strong positive associations between V_cmax_ and J_max_ across both locations (r = 0.70-0.76, p < 0.0001; Figure 3E), consistent with the coordinated regulation of these biochemical traits. LMA showed a strong association with both V_cmax_ and J_max_ (r = 0.50-0.65, p < 0.0001). There was a weaker, but still significant correlation between SPAD, V_cmax_ and J_max_ (r = 0.16-0.38, p < 0.0001), reflecting the established link between leaf nitrogen allocation and photosynthetic capacity.

### Narrow leaf morphology reduces leaf area index, with effects modulated by row spacing and location

Narrow leaf morphology consistently reduced canopy-level LAI across all treatment combinations, though the magnitude of this effect varied with both row spacing and location. By tracking the progression of LAI across the growing season (Figure 4A-B, S6A-B), we identified distinct patterns in canopy dynamics influenced by all three experimental factors.

**Figure 4.**
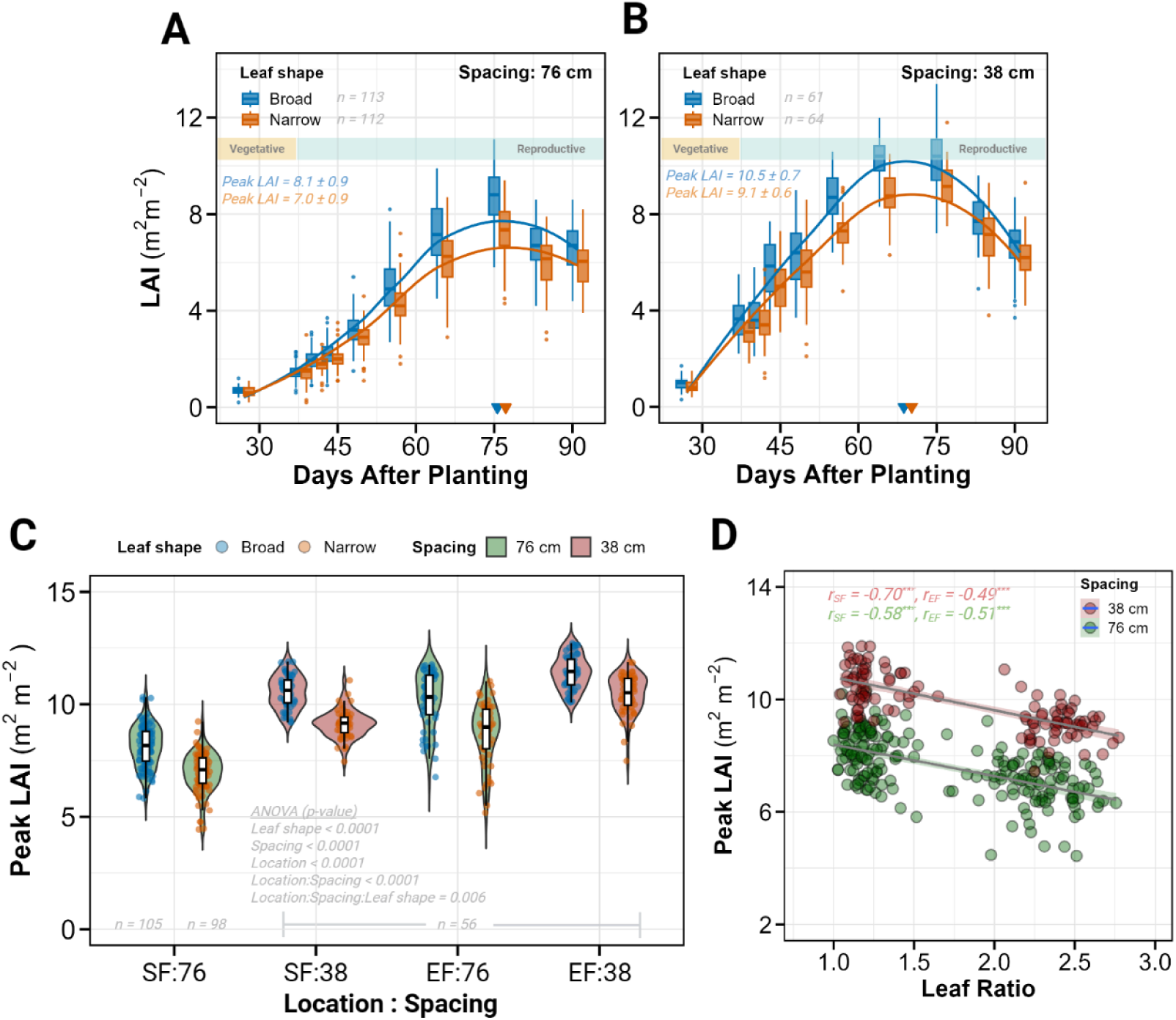
Temporal dynamics and association of Leaf Area Index (LAI). **A-B)** Time course of LAI measurements at SoyFACE under 76-cm (A) and 38-cm (B) row spacing. Box and whisker plots display LAI evolution at ten time points for broad (blue) and narrow (orange) leaf shapes, with fitted growth curves in corresponding colors. The horizontal rectangle at the top indicates vegetative and reproductive growth stages. Colored triangles on the x-axis mark the timing of peak LAI for each leaf shape. Sample sized (n) and peak LAI values (± SD) are provided for both leaf shapes. **C)** Violin plots with embedded box and whisker plots comparing peak LAI across all location-spacing combinations. Plots are colored by row spacing (76-cm in green, 38-cm in red), with individual data points colored by leaf shape (broad in blue, narrow in orange). Sample sizes (n) and ANOVA p-values from mixed model analysis are displayed. **D)** Scatter plot showing the relationship between leaf ratio (x-axis_ and peak LAI (y-axis) for location SF. Data points are colored by row spacing (38-cm in red, 76-cm in green), with corresponding regression lines and shaded 95% confidence intervals. Pearson correlation coefficients (r) are provided for each location (SF, EF) and spacing treatment with significance indicated by asterisks (* p < 0.05, ** p < 0.01, *** p < 0.001).

LAI increased progressively from early vegetative stages (25 days after planting) until reaching maximum values during mid-reproductive development. Peak LAI occurred a week earlier in 38-cm rows (approximately 68 days after planting) than in 76-cm rows (approximately 75 days after planting) across both locations (Figure 4A-B, S6A-B). Within each spacing treatment, narrow-leaf lines reached peak LAI slightly later than broad-leaf lines (1.5 days later at SoyFACE and 2.5 days later at Energy Farm).

All three experimental factors significantly affected peak LAI with comparable effect sizes (0.51-0.70, p < 0.0001; Figure 1B-C), indicating their similar importance in determining canopy development. Location had a substantial effect, with Energy Farm plots achieving 16% higher peak LAI than SoyFACE plots (10.2 vs. 8.59, p < 0.0001). Row spacing exerted an even stronger influence. The 38-cm spacing rows had 20% higher peak LAI compared to 76-cm spacing rows (10.39 vs. 8.5, p < 0.0001).

Most importantly for our research question, leaf morphology consistently affected peak LAI, with narrow-leaf lines showing 13.3% lower values compared to broad-leaf lines (8.8 vs 10.1, p < 0.0001; Figure 4A-C, S6A-B). This reduction ranged from 8.7% to 15.3% across the four location-spacing combinations, indicating the consistency of this effect despite environmental and management differences. The negative association between leaf shape and canopy development was further supported by significant negative correlations between leaf ratio and peak LAI across all treatments (r = −0.49 to −0.70, p < 0.0001; Figure 4D).

### Row spacing drives canopy coverage dynamics while leaf shape has minimal impact

Despite significant differences in LAI, broad and narrow leaf types achieved similar canopy closure rates, suggesting compensatory mechanisms in canopy development. We analyzed temporal dynamics of canopy coverage (CC) using 20 aerial imaging timepoints spanning from early vegetative development (17 days after planting) to complete canopy closure at the SoyFACE location (Figure 5A-B).

**Figure 5.**
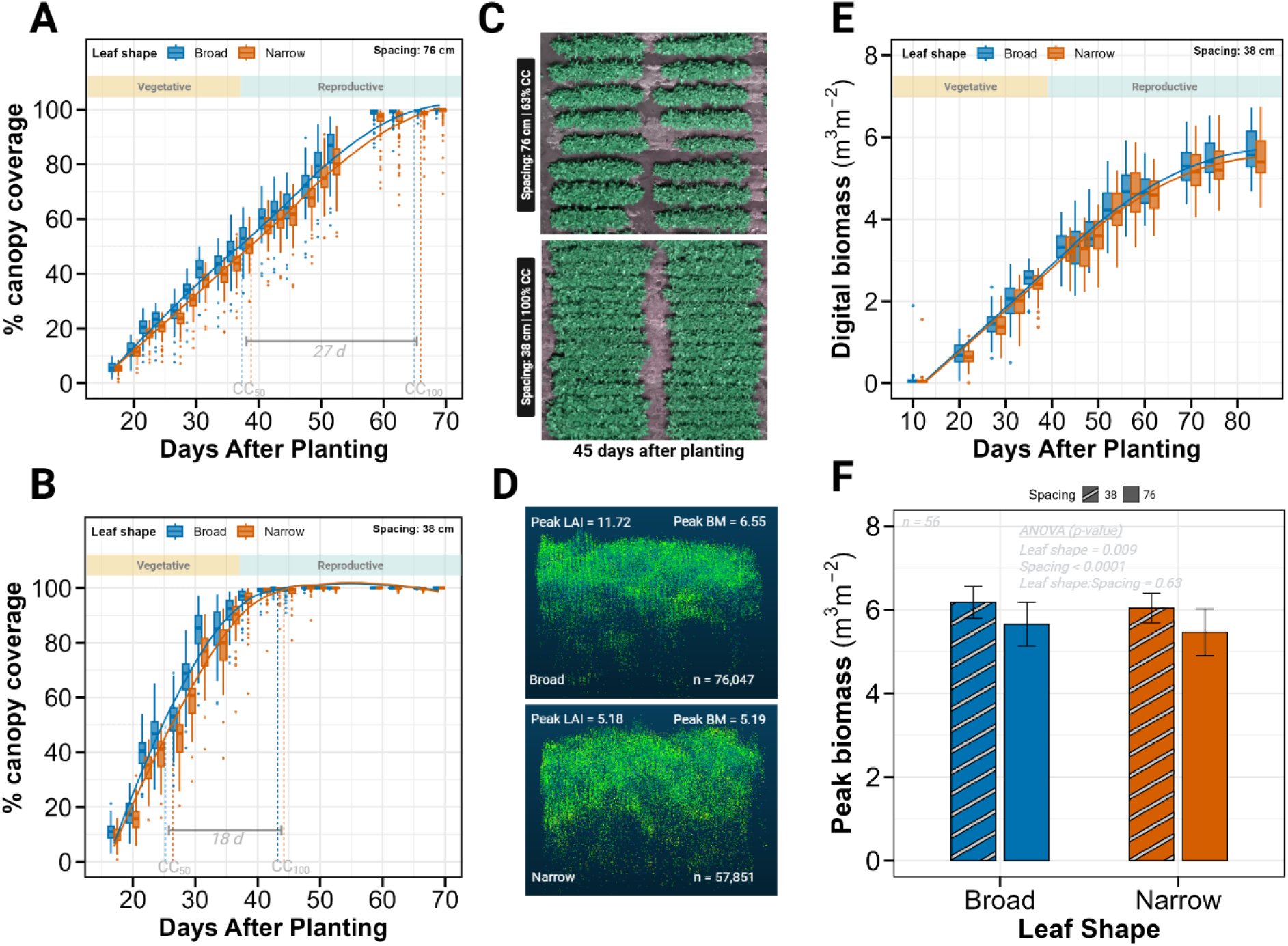
Canopy development dynamics and biomass accumulation across leaf types and row spacings. **A-B)** Temporal progression of percent canopy coverage at SoyFACE for 76-cm (A) and 38-cm (B) row spacing. Box and whisker plots show distributions at 20 time-points for broad (blue) and narrow (orange) leaf shapes, with fitted curves in corresponding colors. The horizontal rectangle indicates vegetative and reproductive growth stages. Vertical colored lines mark the days to reach 50% (CC_50_) and 100% (CC_100_) canopy coverage for each leaf type, with horizontal bars indicating the time interval between these thresholds. Data were derived from drone imagery. **C)** Aerial imagery of experimental plots 45 days after planting, showing 76-cm spacing (upper panel) and 38-cm spacing (lower panel). The image was captured when 38-cm plots approached 100% canopy coverage while 76-cm plots reached approximately 60% coverage. **D)** Three-dimensional point cloud visualizations (visualized in two dimensions) from representative broad leaf (upper panel) and narrow leaf (lower panel) plots at Energy Farm. Point clouds are displayed in green against a dark blue background. Peak LAI values, peak biomass values, leaf shape classification, and number of point clouds (n) are indicated for each plot. **E)** Time course of digital biomass accumulation at Energy Farm under 76-cm spacing. Box and whisker plots show distributions at 14 time points, with the horizontal rectangle indicating vegetative and reproductive growth stages. **F)** Peak biomass comparison between broad (blue) and narrow (orange) leaf shapes across both spacings. Bars with white stripes represent 38-cm spacing, while solid bars represent 76-cm spacing. Error bars indicate standard deviation. Sample size (n) and ANOVA p-values are provided. Digital biomass data was collected using LiDAR attached to cable-mounted Spidercam^TM^ platform.

Row spacing was as the dominant factor controlling the time to canopy closure, with large significant effects on both days to 50% canopy coverage (CC_50_; effect size = 1.81, p < 0.0001) and days to complete canopy closure (CC_100_; effect size = 2.0, p < 0.0001; Figure 1B-C). Plots with 38-cm spacing reached CC_50_ and CC_100_ substantially earlier than those with 76-cm spacing (CC_50_: 26 vs. 38 days, CC_100_: 44 vs. 65 days after planting; Figure 5A-C). The period between half and complete canopy closure was also shorter for 38-cm rows (18 days) compared to 76-cm rows (27 days).

In contrast to the strong effect of row spacing on canopy closure, leaf shape had statistically significant but small effects (CC_50_: effect size = 0.17, p = 0.01; CC_100_: effect size = 0.09, p = 0.007; Figure 1B-C). The practical differences between leaf types were minimal, with narrow-leaf lines reaching CC_50_ and CC_100_ only 0.04 to 1.46 days later than broad-leaf lines (p < 0.001). This negligible delay in canopy closure occurred despite the substantial reduction in LAI observed in narrow-leaf lines.

### Biomass accumulation responds strongly to row spacing with small effects of leaf morphology

We assessed canopy-level biomass dynamics to determine whether the LAI differences between leaf types translated into significant changes in biomass accumulation. A proxy for digital biomass (BM, m^3^ m^-2^) measurements, which quantify three-dimensional canopy volume, was recorded at 14 timepoints throughout the growing season at the Energy Farm location (Figure 5D-F, S6C).

Row spacing was the predominant factor affecting peak biomass accumulation, with an effect size four times larger than that of leaf shape (0.96 vs. 0.24, p < 0.0001 and p = 0.16, respectively; Figure 1B-C). Plots with 38-cm spacing achieved 10% higher peak biomass compared to 76-cm spacing (6.11 vs. 5.56 m^3^ m^-2^, p < 0.0001; Figure 5F), consistent with greater LAI.

Leaf morphology had a smaller but detectable influence on biomass accumulation. Broad-leaf lines produced approximately 3% higher peak BM compared to narrow-leaf lines (p = 0.009; Figure 5F), with this difference ranging from 2% in 38-cm spacing to 3.5% in 76-cm spacing. This modest biomass reduction in narrow-leaf lines was proportionally smaller than the 13.3% LAI reduction (reported in Section 3), suggesting compensatory mechanisms in canopy architecture that partially mitigate the impact of reduced leaf area on overall biomass.

We also analyzed peak Normalized Difference Vegetation Index (NDVI) values from drone imagery at SoyFACE, which provides an integrated measure of canopy greenness and density. Interestingly, neither leaf shape nor spacing significantly affected peak NDVI (effect sizes = 0.1 and 0.96, p = 0.54 and p = 0.70, respectively; Figure 1B-C). Both spacing treatments achieved near identical peak NDVI values (0.938 vs. 0.936 for 38-cm and 76-cm, respectively, p < 0.001), as did both leaf types (0.938 vs. 0.936 for broad and narrow, respectively, p < 0.001; Figure S6D-E). This negligible difference (0.25%) in peak NDVI despite measurable differences in LAI and biomass suggests that NDVI saturates at high canopy density, limiting its sensitivity for detecting subtle differences in fully developed canopies.

### Lines with narrow leaf morphology have altered seed number per pod and maintain grain yield

Despite significant differences in LAI and modest differences in biomass accumulation, narrow-leaf lines maintained yield parity with broad-leaf lines while showing distinctive changes in seed packaging and quality parameters.

Location and row spacing strongly influenced grain yield (effect sizes = 0.75 and 0.91, respectively, p < 0.0001; Figure 1B-C), whereas leaf shape had no significant effect (effect size = 0.08, p = 0.54). Plots at the Energy Farm yielded 17% greater than plots at SoyFACE (6255.9 vs. 5331.6 kg ha^-1^, p < 0.0001), and 38-cm spacing produced 12% greater yield than 76-cm spacing (6108.0 vs. 5449.4 kg ha^-1^, p < 0.0001). Most importantly, the yield difference between broad and narrow leaf isogenic lines was statistically non-significant and agronomically negligible (5801.2 vs. 5756.3 kg ha^-1^, a difference of only 0.78%, p = 0.43; Figure 6A, S9A). This yield parity was consistent across both locations and spacings treatments, demonstrating that narrow-leaf lines can maintain productivity despite reduced leaf area and biomass.

**Figure 6.**
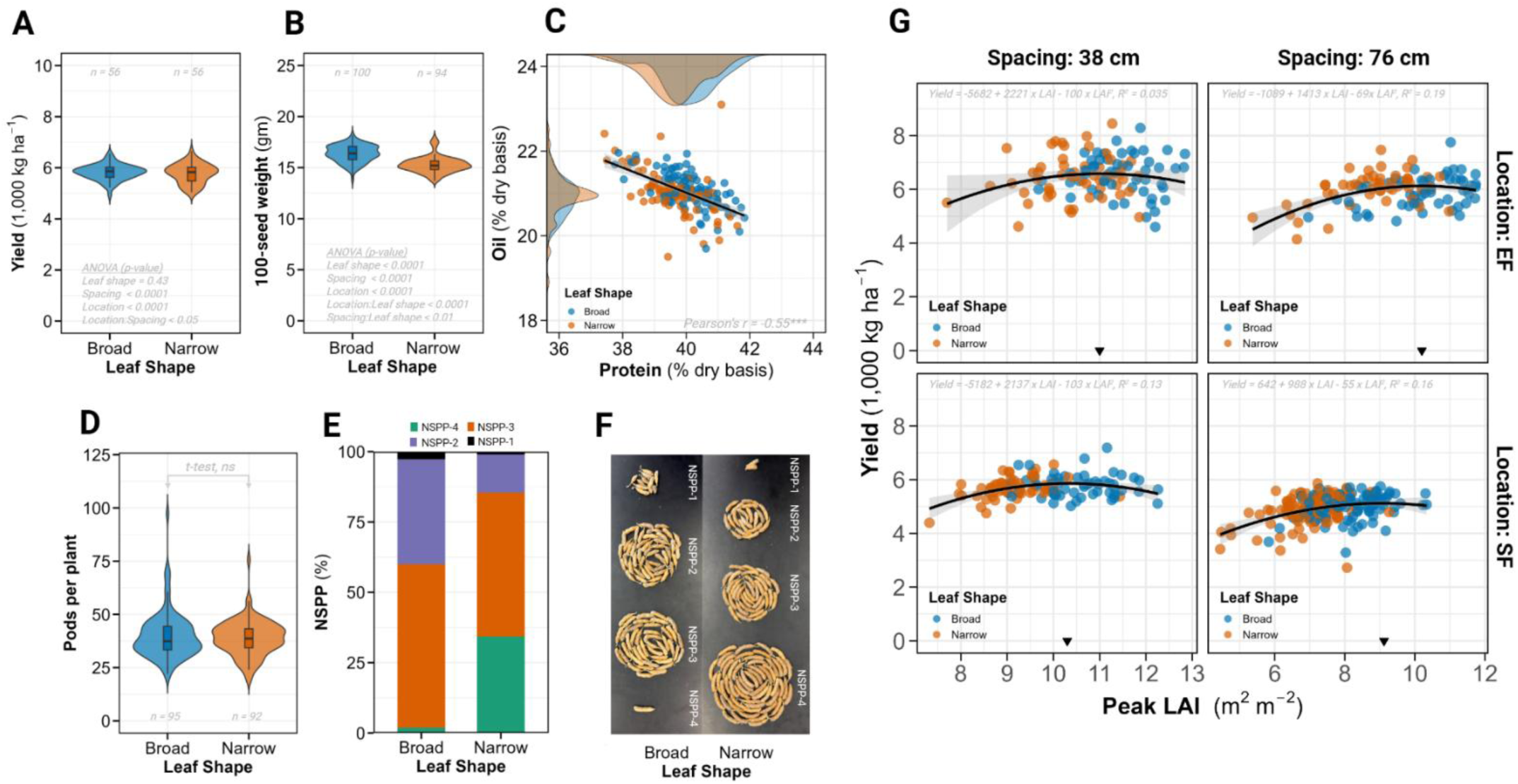
Yield components, seed quality, and yield relationship with LAI. **A)** Violin plot comparing grain yield between broad and narrow leaf shapes, with sample size (n), and ANOVA p-values indicated. Values represent best linear unbiased estimates (BLUEs) from the statistical model. **B)** Violin plot comparing 100-seed weight between broad and narrow leaf shapes. Sample size (n) and ANOVA p-values are provided. Values represent best linear unbiased estimates (BLUEs) from isogenic lines across all locations and spacings. **C)** Scatter plot showing the relationship between seed protein (x-axis) and seed oil content (y-axis), with data points colored by leaf shape. The black regression line includes a shaded 95% confidence interval. Density plots for oil and protein content are displayed along the corresponding axes, colored by leaf shape. Pearson correlation coefficient (r) is provided with significance indicated by asterisks (* p < 0.05, ** p < 0.01, *** p < 0.001). Data includes all locations and spacings. **D)** Violin plot comparing pod number per plant between broad (blue) and narrow (orange) leaf shapes, with sample size (n) and t-test p-value indicated, Data collected only at SoyFACE 76-cm row spacing. **E)** Stacked bar chart showing the percentage distribution of seeds per pod (NSPP) for broad and narrow leaf shapes. Each bar is subdivided to represent the proportion of 4-seeded (NSPP-4, green), 3-seeded (NSPP-3, orange), 2-seeded (NSPP-2, purple) and 1-seeded (NSPP-1, black) pods. Data collected only at SoyFACE 76-cm row spacing. **F)** Representative images of soybean pods categorized by seed number per pod for broad leaf (left panel) and narrow leaf (right panel) shapes. **G)** Quadratic relationship between peak LAI (x-axis) and grain yield (y-axis) for each location and spacing combination. Data points are colored by leaf shape, with black triangles indicating the optimal peak LAI value that maximizes yield. Black curves represent quadratic regression fits with shaded confidence intervals. The quadratic equations and coefficient of determination (R^2^) for each model are provided.

While yield was unchanged, 100-seed weight (SW) was significantly affected by all three factors, with leaf shape exerting the largest effect (effect size = 1.43, p < 0.0001; Figure 1B-C). Seeds from broad-leaf lines were 9% heavier than those from narrow-leaf lines (16.49 vs. 15.01 g per 100 seeds, p < 0.0001; Figure 6B). Location and spacing also influenced SW, with Energy Farm producing 5% heavier seeds than SoyFACE (16.2 vs. 15.4 g, p < 0.0001) and 38-cm spacing yielding 2% heavier seeds than 76-cm spacing (15.96 vs. 15.64 g, p < 0.0001).

Seed composition analysis revealed small but significant differences in protein content, with leaf shape showing a four-fold larger effect (effect size = 0.62, p < 0.001) than location or spacing (effect sizes = 0.16 and 0.19, p = 0.29 and p = 0.20, respectively, Figure 1B-C). Broad-leaf lines had 1.4% higher protein content than narrow-leaf lines (40.32% vs. 39.76%, p < 0.0001). Conversely, oil content showed minimal variation between leaf types (20.99% vs. 20.95%, p = 0.66) but varied significantly by location and spacing (effect sizes = 0.85 and 0.73, p < 0.0001). The inverse relationship between protein and oil was apparent in the narrow and broad leaf lines (r = −0.55, p < 0.0001; Figure 6C).

The most striking difference between leaf types appeared in seed distribution patterns. Analysis of pod characteristics from SoyFACE plots with 76-cm spacing revealed dramatic differences in the number of seeds per pod (NSPP) despite no significant differences in total pod number per plant (40 vs. 39 for broad vs. narrow, effect size = 0.11, p = 0.44, Figure 1B-C, 6D). Narrow-leaf lines had many four-seeded pods, with NSPP-4 accounting for 34% of total pods compared to only 1.8% in broad-leaf lines (effect size = 1.72, p < 0.0001; Figure 6E-F). Corresponding reductions occurred in three-seeded pods (51.2% vs. 58.2%), two-seeded pods (13.6% vs. 37.3%), and one-seeded pods (1% vs. 2.7%) in narrow vs. broad leaf lines (p < 0.001). This increase in four-seeded pods in narrow-leaf lines effectively compensated for their 9% reduction in individual seed weight, resulting in the observed yield parity between leaf types.

### Canopy optimization for yield follows a nonlinear relationship with peak LAI

To understand the functional relationship between canopy development and productivity, we analyzed the mathematical relationship between peak LAI and grain yield across all treatment combinations. When comparing linear versus quadratic models of the LAI-yield relationship, the quadratic model consistently provided superior fit across all location-spacing combinations. Model comparison metrics strongly favored the quadratic relationship, with lower AIC values (3593 cs. 3616), generally lower BIC values (3637 vs. 3651), reduced error (RMSE: 31.8 vs. 32.9), and higher variance explanation (R^2^: 0.52 vs. 0.28) cumulatively across all conditions (Figure S9B). This indicated that yield does not increase linearly with increasing LAI but rather reaches a maximum at an optimal LAI value before declining.

The estimated optimal LAI values for maximizing yield varied by treatment combination, ranging from 9.06 (SoyFACE, 76-cm spacing) t0 11.0 (Energy Farm, 38-cm spacing). The intermediate values were remarkably similar for the remaining conditions: 10.2 for Energy Farm with 76-cm spacing and 10.3 for SoyFACE with 38-cm spacing (Figure 6G). The moderately low R^2^ values of these quadratic models (ranging from 0.035 to 0.19) indicate that while LAI is an important determinant of yield, substantial yield variability is explained by other factors.

### Multivariate analysis of leaf traits and canopy architecture

While the quadratic relationship between LAI and yield provided insight into canopy optimization, it represented only one component of the complex trait network determining soybean productivity. Therefore, to synthesize the complex relationships among the leaf morphological, canopy architecture and yield components traits, two complementary multivariate approaches were employed: Principal Component Analysis (PCA) and Random Forest (RF) regression. PERMANOVA analysis of the multivariate data structure revealed that while leaf shape, spacing and location had statistically significant effects on trait relationships, the interaction effects explained minimal variation (< 0.4%), supporting a global approach to multivariate analysis rather than treatment-specific analyses.

#### PCA reveals coordinated variation in leaf and canopy traits

PCA identified two primary axes of variation that together explained 66.1% of the total variance in trait relationships. The first principal component (PC1), accounting for 46.1% of the variance, was strongly associated with leaf morphological and LAI traits (loadings > 0.8) (Figure 7A). Notably, photosynthetic traits (V_cmax_ and J_max_) and LMA were negatively correlated with PC1, demonstrating a physiological trade-off with leaf structural traits. PC2 explained an additional 20% of the variance and was primarily defined by LR (loading = 0.85) and leaf length (loading = 0.77), suggesting a separate axis of variation for leaf shape independent of size. When individual lines were plotted along these components, broad-leaved and narrow-leaved isogenic lines separated primarily along PC2, with minimal overlap between groups (Figure 7B). This separation confirms that our introduced genetic variation primarily affected leaf shape with limited pleiotropic effects of other traits.

**Figure 7.**
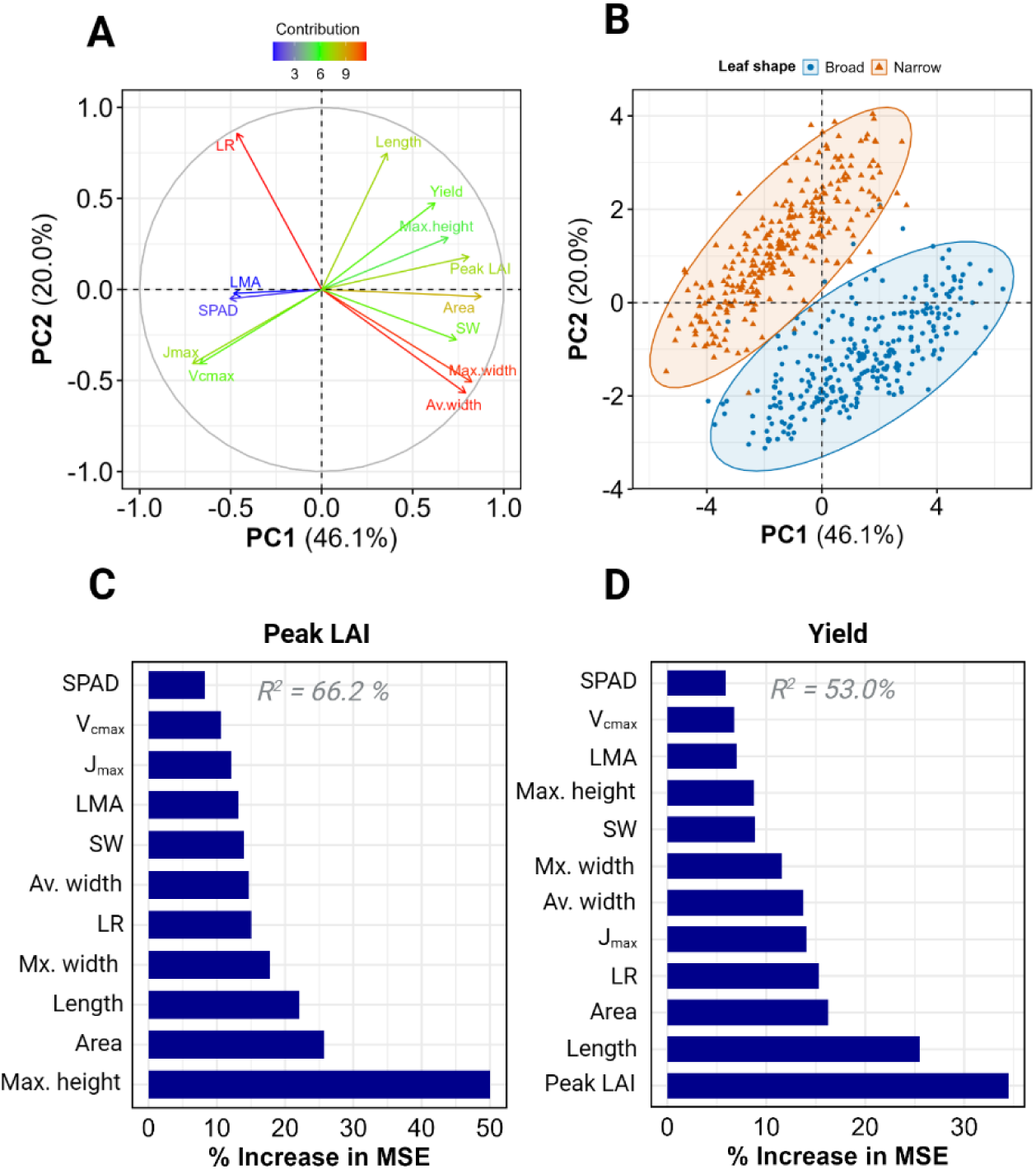
Multivariate analysis of leaf shape effects on yield and related traits. **A)** PCA loading plot displaying the contribution and directionality of individual variables to PC1 and PC2. Arrows represent variables colored according to their contribution magnitude, ranging from highest (red) to lowest (blue) with intermediate values in green. **B)** Principal component analysis (PCA) of phenotypic traits showing PC1 and PC2, which collectively explain the primary variance in the dataset. Individual data points represent lines from all locations and spacings, colored by leaf shape (broad: blue circles, narrow: orange triangles). Two larger central point indicate group centroids, with shaded ellipses demarcating the distribution of each leaf shape category. **C-D)** Random Forest variable importance plots for Peak LAI (C) and Yield (D). The x-axis displays the precent increase in mean square error (MSE) when each predictor variable (y-axis) is randomly permuted, indicating the variable’s predictive importance. Model fit (R^2^) values are provided with each panel.

#### RF analysis ranks the relative importance of traits for predicting yield and peak LAI

RF regression models were constructed to identify key predictors of peak LAI and grain yield, with the models explaining 66.2% and 53.0% of the variance, respectively. For peak LAI, plant height was the strongest predictor (50.0% increase in MSE), followed by leaf area (25.7%) and leaf length (22.1%) (Figure 7C). Interestingly, leaf ratio ranked relatively lower in importance for peak LAI prediction (15.1%), although leaf ratio is correlated with both leaf area and leaf length. For grain yield prediction, peak LAI emerged as the most important predictor variable, with a 34.5% increase in MSE, followed by leaf length (25.5%), leaf area (16.2%) and leaf ratio (15.3%) (Figure 7D). This hierarchy underscores the primary importance of canopy development for determining yield potential, with leaf morphological traits playing significant but secondary roles.

## Discussion

Agricultural productivity hinges on balancing resource investment. The Green Revolution of the 1960’s significantly boosted harvest indices in crops through optimized resource use (Miflin, 2000). However, modern crops face new challenges in resource allocation. Artificial selection in modern varieties has indirectly led to maximizing leaf surface area (reviewed in Slattery and Ort, 2021). Combined with rising atmospheric CO_2_ concentrations, this trend can result in overinvestment in leaf development, diverting resources from reproductive growth to vegetative structures (Srinivasan et al., 2017; Ainsworth and Long, 2005; Soares et al., 2021). To address this, we developed isogenic soybean lines with reduced leaf area by introducing narrow leaf morphology into a broad leaf elite modern variety. Evaluating these lines across two distinct environments and agronomically relevant row spacings enabled us to assess how reduced leaf investment influences canopy structure, physiology and yield outcomes.

### Leaf shape optimization balances resource allocation

Narrow leaf morphology in soybeans improved resource allocation, maintaining yield despite a 13% decrease in leaf area index (LAI) and 3% reduction in biomass (Fig. 2C, 5F, 6A). Narrow leaf shape did not change yield (< 1% difference, p = 0.43) across environments and row spacings. These results align with prior studies that compared productivity of narrow and broad leaf soybeans (Mandl and Buss, 1981; Wells et al., 1993; Dinkins et al., 2002). In contrast, Cai et al. (2021) reported a ∼9% yield increase in narrow leaf mutants of a low-latitude cultivar in China while Bianchi et al. (2020) found yield benefit only under high-density planting in Argentina, reflecting important genotype x environment interactions. Our study of over 200 BC_3_F_2:3_ lines suggest that reducing vegetative investment may improve the harvest index and promote yield efficiency in soybeans.

The maintenance of yield despite the reduced leaf area implies that narrow-leaf lines achieve greater return on structural investment. Bianchi et al. (2020) demonstrated that narrow-leaf lines enhance light penetration to lower canopy layers, reducing self-shading and enhancing photosynthesis. Additionally, Tamang et al. (2023) reported that those narrow leaves, with a higher leaf length-to-width ratio, are thicker and possesses more spongy mesophyll, which potentially increases light scattering and absorption. Our findings support this, showing a modest (∼4%) increase J_max_ in narrow-leaf lines (p < 0.05), with V_cmax_ unaffected (p > 0.2, Fig. 3D). While photosynthetic biochemistry remained largely unchanged amongst broad and narrow lines, a 5% increase in LMA in narrow leaf lines suggests increased construction costs compensated by enhanced carbon gain and longevity (Poorter et al., 2009). The strong correlation between LMA and both V_cmax_ and J_max_ (Fig. 3E) reinforces the role of LMA in photosynthetic capacity, consistent with optimal resource allocation theory (Bloom et al., 1985), which posits that plants distribute resources to optimize fitness under constraints. In agricultural contexts, where reproductive output (yield) is the primary fitness metric, the narrow-leaf trait may help optimize the trade-off between vegetative and reproductive investment.

### Narrow leaf morphology maintains canopy coverage likely through architectural compensation

Early canopy coverage is critical in agricultural crops, as it enhances light interception, suppresses weed growth, and maximizes photosynthetic productivity during key early vegetative growth stages (Richards, 2000; Purcell et al., 2002). Despite a 13% reduction in peak LAI (Fig. 4C), our narrow leaf isogenic soybean lines showed no difference in canopy development dynamics. The negligible delay in canopy closure (less than 2 days delay to achieve 100% canopy closure, Fig. 5A, B) and only modest reductions in biomass accumulation (3% lower at high yielding location Energy Farm, Fig. 5F) suggest architectural adaptations that partially mitigate the impact of reduced leaf area. This compensation likely involves alterations in leaf arrangement, canopy structure, and light interception patterns that collectively maintain effective resource capture despite the modified leaf morphology (Niinemets, 2010; Sreekanta et al., 2024). While LAI and biomass differed significantly, NDVI did not vary between leaf types or spacings at peak canopy size (Fig. S6D, E). This saturation of NDVI under high-density canopies emphasizes the limitations of traditional greenness indices for distinguishing subtle morphological effects and suggests the need for integrating structural or multispectral indices in high-resolution phenotyping.

### Nonlinear relationship between LAI and yield

We observed a nonlinear (quadratic) relationship between peak LAI and grain yield, where yield peaked at an optimal LAI and declined thereafter (Fig. 6G). This optimal LAI varied with leaf type, location and spacing, ranging from 9 (SoyFACE, 76-cm spacing, narrow leaves) to 11 (Energy Farm, 38-cm spacing, broad leaves). These values exceed earlier thresholds (e.g. LAI = 7.5 in Srinivasan et al., 2017), likely due to differences in genotype, planting density, management, and environmental conditions.

This nonlinear relationship between canopy size and seed yield supports the hypothesis that LAI in soybean is too high (Srinivasan et al., 2017). Beyond the optimal LAI, excess foliage may increase respiration, mutual shading or create microclimate stress, reducing reproductive output (Board and Harville, 1992; Setiyono et al. 2008; Tagliapietra et al., 2018; Digrado et al., 2020). Narrow leaf lines may inherently position plants closer to this optimum, achieving high yield with reduced vegetative cost. Supporting this, our Random Forest regression identified peak LAI as the most important trait predicting grain yield, followed by leaf architectural traits (Fig. 7D). These results suggest that optimizing whole-plant architecture has a stronger influence on yield than leaf shape alone, highlighting the importance of integrated phenotypic strategies in soybean ideotype design.

### Environmental context-dependency of leaf shape effects

The adaptive value of architectural traits, including leaf morphology, is highly contingent on environmental conditions (Sarlikioti et al., 2011). Krause et al. (2025) highlighted the importance of environmental context by defining soybean mega-environments. Our study underscores this, with location significantly influencing traits such as V_cmax_, J_max_, peak LAI, and yield (Fig. 1B, 3C, D). Energy Farm, the higher productivity site had 17% higher grain yield, 16% higher peak LAI, and 5% heavier seeds than SoyFACE (Fig. 4C, 6A, B).

Interestingly, V_cmax_ and J_max_ were significantly higher at SoyFACE (17% and 12%, respectively, Fig. 3C, D), indicating that in resource-limited environments, soybeans increase photosynthetic capacity per unit leaf area to compensate for lower canopy development. We suspect that greater fertility at the Energy Farm enabled greater total leaf area (∼44% more), allowing nitrogen to be distributed across more tissue. This led to slightly lower nitrogen concentration per leaf, evidenced by 3% lower SPAD readings, and correspondingly lower photosynthetic capacity (Evans, 1989; Poorter and Evans, 1998; Walker et al., 2014). These results support the hypothesis that elevated photosynthetic capacity is particularly important in stressful environments, a key insight for breeding resilient soybean varieties (Flood et al., 2011).

### Row spacing effects on canopy architecture and yield

Row spacing strongly influences resource competition by altering plant spatial distribution (Board et al., 1990; Andrade et al., 2002). Narrower rows promote faster canopy closure and greater light capture, and suppress weed density, biomass and weed seed production (Singh et al., 2023). In our study, canopy closure was 3 weeks earlier in narrow rows, associated with a 20% increase in LAI (Fig. 4C), 10% increase in biomass (Fig. 5F) and 12% yield boost (Fig. 6A). These benefits were consistent across leaf types, suggesting that optimized morphology and management practices can be combined to enhance resource use efficiency. This supports broader findings by Andrade et al. (2019), who documented up to 18% yield increases in narrow rows across nearly 5,000 US soybean fields, especially under late planting or short-season conditions. A trade-off between narrow row spacing and disease pressure has also been reported (Webster et al., 2022), so the economic gains of greater yields in narrow row spacing would need to be considered against greater costs of fungicides by growers.

### Ecological context and breeding implications

Leaf morphology represents a fundamental axis of plant functional diversity that influences light capture, water relations, gas exchange and herbivore defense across ecosystems. Our findings partially align with leaf economic spectrum (LES; Wright et al., 2004) theory – narrow leaf lines invest in higher LMA with reduced total leaf area, signaling conservative resource-use strategies. However, they deviate by maintaining yield despite this reduced leaf area, primarily through increased reproductive efficiency. Such within-species variations often challenge global leaf economic spectrum patterns, revealing more complex resource allocation strategies than simple trade-offs, particularly in crop species where breeding has selected for novel trait combinations that optimize productivity (John and Garnica-Diaz, 2023).

These results emphasize the need to reassess traditional breeding priorities. Historically, selection for high yield has resulted in broad leaves and high LAI to maximize early-season growth and light interception. However, our study shows that narrow leaf traits can reduce vegetative costs without sacrificing yield, potentially contributing to greater efficiency under both high-input and stress-prone environments. Given increasing climate variability and input limitations, this shift in breeding priorities could enhance crop resilience.

The narrow-leaf phenotype is largely controlled by a single gene, *GmJAG1* (Jeong et al., 2012), which simplifies introgression into elite backgrounds. Previous work has shown minimal pleiotropic effects, which supports its utility in breeding pipelines. This is further reinforced by PCA analysis, which showed clear separation of leaf types along leaf shape axes while other traits remained largely unchanged (Fig. 7B), validating the value of these near-isogenic lines for physiological studies.

The near-isogenic lines developed here, which differ mainly in leaf morphology, provide valuable materials for physiological modeling. They offer unique opportunities to isolate the effects of LAI and canopy structure on productivity in a controlled genetic background, enabling more precise parameterization of crop growth models and supporting efforts to simulate genotype x environment x management interactions.

### Future directions

Our field study was done in East Central Illinois, in the heart of the U.S. Corn Belt. Future research should evaluate the performance of narrow leaf genotypes in a broader spectrum of environmental conditions, including regions prone to water limitation, high temperatures, or suboptimal planting densities. Such conditions may accentuate the advantages of reduced leaf area, improving water use efficiency and heat dissipation while minimizing excessive vegetative growth. In addition, mechanistic studies on carbon partitioning and source-sink dynamics in narrow leaf lines would provide insights into how these genotypes maintain yield. Investigating the interaction between leaf shape and other architectural traits- such as leaf angle, plant height, branching architecture, and rooting depth- could help refine ideotype models tailored to different agroecological zones.

From a technological perspective, integrating narrow leaf morphology with novel plant protection strategies could improve performance in places with high disease load. Additionally, the narrow leaf trait could be combined with strategies to enhance water use efficiency, abiotic stress resistance, photosynthetic, sink strength or nutrient uptake to enhance performance in suboptimal environments (Bailey-Serres et al., 2019). Fast-tracking combinations of strategic traits could enhance the development of climate-resilient, resource-efficient soybean varieties tailored for sustainable production in a changing world.

## Conclusion

Based on the analysis of 204 isogenic soybean lines, this study demonstrates that narrow leaf morphology is a promising strategy for optimizing resource allocation without compromising grain yield. Despite reducing LAI by 13% and biomass by 3%, narrow-leaf lines maintained yield parity with broad-leaf counterparts across different environmental conditions and management practices, achieving greater return on structural investment. The narrow-leaf lines had similar canopy closure dynamics and photosynthetic capacity, but altered seed packaging patterns with significantly more four-seeded pods. The nonlinear relationship between peak LAI and yield supports the hypothesis that modern soybean varieties may overinvest in vegetative structures at the expense of reproductive efficiency. These findings challenge traditional breeding paradigms that have resulted in high leaf area and suggest that the single-gene control of narrow leaf morphology through *GmJAG1* offers a tractable approach for developing resource-use efficient cultivars that maintain productivity while reducing metabolic costs associated with excessive canopy development. We offer a new strategy to test for increasing sustainable agriculture under intensifying climate variability and resource constraints.

## Materials and Methods

### Parent selection and development of isogenic lines via repeated backcrossing

To develop isogenic soybean lines that varied for leaf morphology, we selected the broad-leaved cultivar LD11-2170 (Maturity Group III, MG III) as the recurrent parent. Developed at the University of Illinois, LD11-2170 has served as a reference check in the Uniform Soybean Tests of the Northern Region since 2019, owing to its consistent high field trial yields (> 4,700 kg per hectare) and favorable agronomic traits (Cai and Brock, 2023). For the narrow-leaf trait, two donor lines PI 612713A (MG I) and PI 547745 (MG II) were identified from a 2020 leaf shape survey of the USDA germplasm collection conducted in Urbana, IL (Tamang et al., 2023). PI 612713A originated from Heilongjiang Sheng, China and was deposited in USDA National Plant Germplasm System (NPGS) in 1999. Similarly, PI 547745 was developed at IL and deposited in NPGS in 1975 (npgsweb.ars-grin.gov). These recurrent and donor lines were further characterized in summer 2022, during which the known leaf shape gene *GmJAG1* (Jeong et al., 2012) was sequenced to confirm the presence or absence of its allele.

Initial crosses to produce F_1_ hybrids were performed in summer 2021 at the University of Illinois. Three successive backcrosses were subsequently conducted to introgress the narrow-leaf trait into the LD11-2170 background from each donor line. During the backcrossing, Kompetitive Allele-Specific PCR (KASP) markers developed from the *GmJAG1* gene were used to select F_1_ plants that were heterozygous for this gene. From the resulting BC_3_F_1_ generation, 600 seeds derived equally from the two narrow-leaved donors were planted. Segregating BC_3_F_2:3_ lines were screened with the KASP marker, yielding 209 homozygous lines: 106 broad-leaved and 103 narrow-leaved (116 from the PI 547745 parent and 93 from the PI 612713A parent).

The BC_3_F_2_ plants were grown and 204 BC_4_F_2:3_ lines were advanced for seed increase. Seeds from these lines (BC_3_F_2:4_) were subsequently planted in experimental plots for this study. The three parental lines (LD11-2170, PI 612713A and PI 547745) and the 204 BC_3_-derived lines were genotyped using the Agriplex 1K Soy SNP panel, comprising 1327 single nucleotide polymorphisms (SNPs) (Agriplex Genomics, Cleveland, OH, USA).

### Field experimental design, planting and management practices

The experimental plots were arranged in a Type II Modified Augmented Design (MAD) (Lin and Poushinsky, 1985). This implementation of the Type II MAD specifically addresses the rectangular nature of soybean field plots by arranging plots in rows within blocks. As described by Lin and Poushinsky (1985), this arrangement ensures optimal spatial control while accommodating row-based planting systems. Each block contained 25 plots with the primary check (LD11-2170) positioned centrally to maintain equal distances to surrounding test plots, maximizing the effectiveness of spatial adjustment. Secondary checks (PI 612713A and PI 547745) were randomly distributed within blocks to capture additional spatial variation that might not be accounted for by the primary check alone.

Field experiments were conducted at two experiment farms, SoyFACE (SF) and Energy Farm (EF) in two row spacings (38-cm and 76-cm), at the University of Illinois, Urbana-Champaign, USA, approximately 2 miles apart. Of the 204 BC_3_F_2:4_ lines, a set of 112 (56 broad-leaved and 56 narrow-leaved isogenic lines) were planted at both locations and spacings. The remaining second set of 92 BC_3_F_2:4_ lines were planted at SoyFACE with a 76-cm spacing. At SoyFACE, this resulted in 9 blocks for 76-cm spacing and 5-blocks for 38-cm spacing. At Energy Farm, 10 blocks were established for each spacing. Each block contained 25 plots, with isogenic lines randomly assigned to achieve approximately equal numbers of broad- and narrow-leaved lines. Across both locations and spacings, a total of 600 plots were planted. Each plot consisted of four 2.4 m rows, separated by 0.9 m alleys. At both locations, the two experimental plots (plots planted at 38-cm and 76-cm row spacing) were surrounded by four border rows of a commercial soybean variety on all sides. Seeds were mechanically planted at 314,000 seeds per hectare for 76-cm spacing and 447,000 seeds per hectare for 38-cm spacing. Planting occurred on May 30, 2024 at Energy Farm and June 4, 2024 at SoyFACE.

Weeds were manually removed as needed. Irrigation was applied once at SoyFACE and twice at Energy Farm during the early vegetative growth stages.

## Data Collection

### Leaf morphology and Leaf Area Index

Leaf morphology was characterized by non-destructively measuring total leaf area, length along the midrib, and widths (average and maximum) using a portable leaf area meter (LI-3000C; LI-COR Biosciences, Lincoln, NE, USA). Leaf Ratio (LR), defined as the ratio of length to maximum width (Chen and Nelson, 2004), was calculated from these measurements. Data was collected once from the middle trifoliate, fully expanded leaf between the 5^th^ and 7^th^ nodes. Leaves destructively harvested for additional measurements (e.g., A-Ci response curves, see below) were also scanned with the leaf area meter to create paired datasets with those variables, as leaf ratios varied slightly by node. Relative greenness of all samples leaves was assessed using a SPAD meter (SPAD-502; Konica Minolta, Tokyo, Japan). These leaves were then dried at 65°C for one week, and their dry mass was recorded to calculate Leaf Mass per Area (LMA).

Leaf Area Index (LAI) was measured throughout the season using the SS1 Canopy Analysis System (Delta-T Devices, Cambridge, UK), a 1-m probe equipped with 64 photosynthetically active radiation (PAR) sensors. The probe was placed perpendicular to the four rows of each plot at the plant base. Ten LAI measurements were taken per plot across all 600 plots at both locations (SoyFACE and Energy Farm), totaling 6,000 measurements, from early vegetative to late reproductive stages (25 to 95 days after planting).

### A-C_i_ response curve measurements

Photosynthetic efficiency of the isogenic lines was evaluated by measuring photosynthetic rates (A) in response to varying intercellular CO_2_ concentrations (C_i_), following established protocols (Long and Bernacchi, 2003; Busch et al., 2024). Measurements were conducted using a portable infrared gas exchange analyzer (LI-6800; LI-COR Biosciences, Lincoln, NE, USA) on leaf samples from 264 plots (135 from SoyFACE, 129 from Energy Farm), representing 127 broad-leaved and 137-narrow-leaved lines. Samples were collected at the R1-R2 developmental stage (Fehr et al., 1971), except for the early-maturing narrow-leaved donor parents (PI 612713A and PI 547745), which were at R3. An undamaged, unshaded, recently matured trifoliate leaf (typically the fourth from the top node) from the middle rows of each plot was tagged one day prior to measurement. The following predawn, tagged leaves were cut at the petiole base, immediately submerged in water, and stored in a cool, dark cabinet to maintain turgor and prevent photosynthetic initiation. Before measurements, leaves were exposed to low light for at least 30 minutes. The LI-6800 was equipped with a 6 cm^2^ circular cuvette, which was clamped onto the middle section of each leaf to accommodate narrower morphologies. Steady-state photosynthetic assimilation rates were recorded at 14 reference CO_2_ concentrations: 400, 300, 200, 100, 75, 50, 25, 400, 400, 600, 800, 1000, 1200 and 1800 ppm. A light intensity of 2000 µmol m⁻² s⁻¹, leaf temperature of 28°C, relative humidity of 65%, fan speed of 10,000 rpm, and airflow of 700 µmol s⁻¹ were maintained in the cuvette.

Data was collected over four days using 9 infrared gas analyzers, with samples grouped by location and row spacing. At the Energy Farm, leaves from 76-cm row spacing plots (n = 62) were measured on July 23, 2024, followed by 38-cm row spacing plots (n = 67) on July 24, 2024 (54-55 days after planting). At SoyFACE, leaves from 76-cm row spacing plots (n = 74) were measured on July 29, 2024, and 38-cm row spacing plots (n = 61) on July 30, 2024 (55-56 days after planting).

### Canopy Coverage, NDVI and Digital Biomass

Canopy coverage of the isogenic lines was quantified using aerial images captured by a DJI Mavic 3 Multispectral Enterprise drone (DJI, Shenzhen, China). Images (captured from an altitude of 29 m with front overlap of 80% and side overlap of 90%) were collected twice weekly between 17 and 99 days after planting (26 flights). Orthomosaics were generated from these images using Pix4D Fields (version 2.8.0; Pix4D, 2024). Canopy coverage was calculated from the red (R), green (G), and blue (B) channels in QGIS (version 3.36; QGIS Development Team, 2024) through a three-step process. First, pixels were classified as soil or vegetation using the Excess Green Index (2G – (R+B)), followed by thresholding to distinguish between the two. Second, a square boundary layer was defined for the middle two rows of each plot, and zonal statistics were used to count total pixels representing vegetation or soil within the boundary. Finally, canopy coverage was expressed as the percentage of vegetation pixels.

Normalized Difference in Vegetation Index (NDVI), an indicator of plant health and proxy for vegetation growth and biomass, was calculated from the same images using the near-infrared (NIR) and red (R) channels with the formula (NIR – R)/ (NIR + R) in QGIS (3.36; QGIS Development Team, 2024). A boundary box encompassing all four rows of each plot was created, and zonal statistics provided mean and median NDVI values per plot. Given the high correlation between mean and median NDVI (r = 0.95), mean NDVI was selected for all subsequent analyses. These two variables, Canopy Coverage and NDVI, were captured only at SoyFACE.

Biomass at the canopy scale was non-destructively measured at 14 different timepoints (between 11 and 84 days after planting) using a LiDAR sensor (OUSTER OS0-128, Ouster Inc., San Franscisco, CA, USA) mounted on a Spidercam^TM^ system (Ross Video Limited, Ottawa, Canada) at Energy Farm. The measurement platform was commanded to scan the plots at a constant rate of 450 mm s^-1^ to record a series of LiDAR frames which were algorithmically converted into point clouds. The digital biomass of the plot was determined by computing the plot volume and dividing the volume by the plot area, where the plot volume was computed using voxel grid algorithm from the open-source Point-Cloud Library (Rusu and Cousins, 2011).

### Plant height and developmental stages

Plant height was manually measured from the base of the plant to the uppermost leaf at all locations and row spacings at six time points, from 25 to 95 days after planting. Concurrently, developmental stages were monitored and recorded at six time-intervals throughout the growing season. These observations enabled the determination of key phenological parameters, including the duration of the vegetative growth phase, days to initiation of the reproductive stage or beginning of flower bloom (R1), and days to full physiological maturity (R8).

### Yield, yield components and quality

#### Pod number and number of seeds per pod

Pod number (PN) and number of seeds per pod (NSPP) were assessed exclusively at SoyFACE, within plots established at a 76-cm row spacing. At maturity, a representative plant was selected from each of 190 plots, consisting of 187 isogenic lines and three parental genotypes. For each plant, the total number of pods was manually counted. Pods were subsequently categorized and counted according to the number of seeds per pod (NSPP), specifically distinguishing among one- (NSPP-1), two- (NSPP-2), three- (NSPP-3), and four-seeded (NSPP-4) pods.

#### Grain yield, seed weight, protein and oil content

Grain yield was determined at maturity by mechanically harvesting all the plots at both locations and spacings using an ALMACO R1 Single-Plot Rotary Combine (ALMACO, Nevada, IA, USA). Harvest data, including total plot weight and grain moisture content, were recorded using the combine’s integrated Weight Hopper System. For plots established with 76-cm spacings, the central two rows were harvested, whereas all rows were harvested from plots with 38-cm row spacings. Yield data were standardized to a moisture content of 13%. Additionally, three random subsamples of 100 seeds each were collected from every harvested plot and weighed to determine seed sizes.

Seed protein and oil content (% dry weight basis) were determined on the same samples used for seed weight assessment, using an NIR instrument (DA 7250 NIR analyzer, PerkinElmer, Waltman, MA, USA). The instrument was calibrated using 10 reference samples with known protein and oil content values previously determined through standard wet chemistry methods. Calibration biases were adjusted based on these reference values, while using the manufacturer’s preset calibrations as a baseline.

## Statistical analysis

All data processing, statistical analyses and visualizations were conducted using R (version 4.2.1, R Core Team, 2024). Descriptive statistics were calculated for all variables, including mean, standard deviation and range. Pearson correlation coefficients (r) were computed to assess relationships among variables, with significance established at p < 0.05. For group comparison, independent sample t-tests were employed for two-group comparisons, while Analysis of Variance (ANOVA) with post-hoc Tukey’s Honestly Significance Difference tests were used for multiple group comparisons. Data wrangling was performed using *dplyr* and *tidyr* packages (version 1.1.4; Wickham et al., 2023; version 1.3.1; Wickham, 2023), and most of the plots were generated with the *ggplot2* package (version 3.5.1; Wickham, 2016).

### Estimating parameters from time course measurements of LAI, CC, NDVI, PH and BM

For traits measured over multiple timepoints such as LAI, Canopy Coverage (CC), Normalized Difference in Vegetation Index (NDVI) Plant Height (PH) and Digital Biomass (BM)- key temporal parameters were extracted by evaluating three curve-fitting models (LOWESS; Locally Weighted Scatterplot Smoothing, logistic, and quadratic with second-degree polynomial). Model selection employed a comprehensive approach, evaluating both traditional fit quality metrics and information criteria. Traditional metrics included Mean Absolute Error (MAE), Residual Sum of Squares (RSS), Root Mean Squared Error (RMSE), and the coefficient of determination (R^2^), while information criteria assessment used Akaike Information Criterion (AIC) and Bayesian Information Criterion (BIC) to account for model complexity. For all traits, logistic models failed to converge during information criteria assessment indicating these traits did not follow a sigmoidal pattern in our experimental context. Formal statistical testing (ANOVA with Tukey HSD) of the traditional metrics confirmed significant differences between models. Across all traits, the LOWESS model consistently showed superior performance, with lower MAE, RSS, RMSE, AIC and BIC, and higher R^2^ values. As a result, the following parameters were extracted from the LOWESS fits: Peak LAI, days to 50% canopy coverage (CC_50_), days to 100% canopy coverage (CC_100_), peak digital biomass, peak NDVI, days to start of NDVI decline and maximum plant height.

### Predicting V_cmax_ and J_max_ from A-ci curves

Photosynthetic parameters, maximum rate of Rubisco activity (V_cmax_) and maximum rate of RuBP regeneration (J_max_), were estimated from A-Ci curves using *PhotoGEA* R package (version 1.1.0; Lochocki et al., 2025).

The *PhotoGEA* package implements a whole-curve fitting approach using maximum likelihood regression and derivative-free optimization algorithms. For our analysis, we employed the *fit_c3_aci* function, which fits A-Ci curves using the Farquhar-von-Caemmerer-Berry (FvCB) model for C_3_ plants. Mesophyll conductance was assumed to be infinite, yielding apparent photosynthetic parameters based on intercellular CO_2_ concentration (C_i_). All parameter estimates were normalized to 25 °C using the temperature response functions of Bernacchi et al. (2001).

### Adjustment for spatial variability in trait estimates

To address the inherent field heterogeneity present in the trial and its influence on the estimates of the 25 traits, two distinct spatial adjustment approaches relevant to the Type II Modified Augmented Design were compared (Method I and Method III, Lin and Poushinsky, 1985). The adjustments were made independently across two locations (SoyFACE and Energy Farm) and two row spacings (38-cm and 76-cm).

The first method (Method I) corresponds to the “adjustment by design structure” approach, which uses an ANOVA model to account for row and column effects through the analysis of control plots. The model follows as:

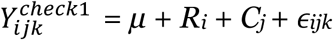

where:

- 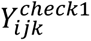 is the observed trait value for the primary check (check1) in row i, column j, and block k,
- *µ* is the overall mean of the trait for the primary check (check1) across all blocks,
- *R_i_ represents* the effect of the *i*-th row,
- *C_j_* represents the effect of the *j*-th column,
- *ϵ_ijk_* is the residual error.

Using the row and column effects estimated from this model, the adjusted trait value for each test plot was computed as:

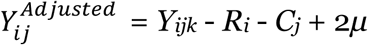

where:

- 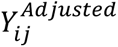 is the adjusted trait value for the test plot in row *i*, column *j*,
- *Y_ijk_* is the observed trait value of the test plot,
- *R_i_* and *C_j_* are the estimated row and column effects,
- *μ* is the overall mean of the primary check.

Similarly, the second method (Method III) corresponds to the “adjustment by regression” approach but extends it by incorporating both primary and secondary checks. The model follows as:

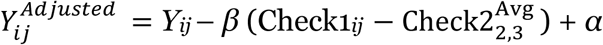

where:

- 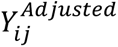 is the adjusted trait value for the test plot,
- *Y_ij_* is the observed trait value,
- Check1*_ij_* is the trait value of the primary check in the corresponding block,
- 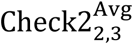 is the average trait value of the two secondary checks,
- *β* is the regression slope relating the secondary checks’ average to the primary check,
- *α* is the regression intercept.

The decision to apply one of the adjustments was based on the relative efficiency (RE) of each approach, defined as:

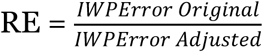

where:

- *IWPError_Original_* is the intra-whole plot error without adjustment,
- *IWPError_Adjusted_* is the intra-whole plot error after applying either method.

The intra-whole plot error was calculated as:

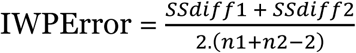

where:

- *SSdiff_1_* and *SSdiff_2_* are the sums of squared differences for the two secondary checks,
- *n_1_ and n_2_* are the number of blocks for the secondary checks.

For the adjusted values, a similar calculation was made for adjusted sums of squared differences (*SSadj_1_* and *SSadj_2_*). The adjustment decision followed these criteria:

- If both methods resulted in RE < 100, no adjustment was applied as precision gains were insufficient.
- If one method had RE ≥ 100 and the other had RE < 100, the method with RE ≥ 100 was chosen.
- If both methods had RE ≥ 100, the method with the higher RE was selected for its greater efficiency in reducing spatial variability.

This comparative strategy was applied individually to each of the 25 traits across the four location and spacing combinations. For each trait and combination, RE values were computed to determine whether adjustment was warranted and, if so, which method to use. Adjustments were not universally applied, as their necessity depended on RE exceeding 100 or the highest RE when both methods met this threshold.

### Best linear unbiased estimates analysis

For yield, seed weight, protein and oil content measured across both locations and spacings, Best Linear Unbiased Estimates (BLUEs) were calculated to obtain representative values for each isogenic line, adjusted for location and spacing effects. BLUEs were generated using linear mixed-effects models implemented in the *lme4* R package (version 1.1.36; Bates et al., 2015), with *emmeans* R package (version 1.10.7; Lenth, 2025) to extract the marginal means (BLUEs) for each line with the following model structure:

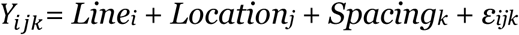

where:

- *Y*_*ijk*_ is the observed trait (grain yield, seed weight, protein or oil content) value,
- *Line_i_* is the fixed effect of the i^th^ isogenic line,
- *Location_j_* is the random effect of the j^th^ location,
- *Spacing_k_* is the random effect of the k^th^ spacing,
- *ε_ijk_* is the residual error.

### Peak LAI-Yield relationship analysis

To investigate the relationship between peak LAI and yield, linear and quadratic regression models were fitted to data for each location (SF, EF) and row spacings (38-cm, 76-cm). Model performance was evaluated using AIC, BIC, R^2^ and Root Mean RMSE to identify the best-fitting model for each group.

### Hierarchical mixed-effects modeling framework to assess trait variation

To evaluate how location, spacing, leaf shape and donor parent affected 25 measured traits, we used hierarchical mixed-effects models implemented in the *lme4* R package (version 1.1.36; Bates et al., 2015). Statistical significance was assessed using the *lmerTest* package (version 3.1.3; Kuznetsova et al., 2017). The preliminary model included four fixed factors: Donor Parent (PI 612713A, PI 547745), Location (SoyFACE, Energy Farm), Spacing (38-cm, 76-cm), and Leaf Shape (Broad, Narrow), along with their interactions and a random intercept for the interaction of Leaf Shape and Isogenic Lines. Since the statistical analysis revealed that Donor Parent did not significantly influence the measured traits (p > 0.05), it was removed from the final model:

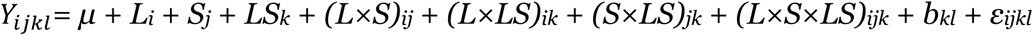

where:

- *Y*_*ijkl*_ is the standardized trait value (z-score),
- *μ* is the overall mean,
- *L_i_* is the fixed effect of Location (SoyFACE, Energy Farm),
- *S_j_* is the fixed effect of Spacing (38-cm, 76-cm),
- *LS_k_* is the fixed effect of Leaf Shape (Broad, Narrow),
- All interaction terms are denoted by products (e.g., (*(L×S)_ij_*),
- *b_kl_* is the random effect for the interaction of Leaf Shape and isogenic line *l*, with 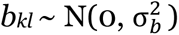, and
- *ε_ijkl_* is the residual error, with 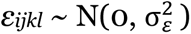.

The models were modified for specific cases based on data type: count data (PN) were analyzed using Poisson or negative binomial GLMMs when overdispersion was detected (overdispersion ratio > 1.5), ratio data (NSPP, LR and NDVI) were handled with beta regression or log transformation, time-to-event data (NDVI decline, CC_50_ and CC_100_) were log-transformed and continuous data were analyzed with linear mixed models. For traits with factors having only one or two levels, the corresponding terms were removed from the model. For cases where convergence issues occurred, the fixed-effects linear model was:

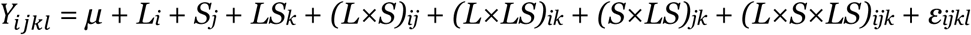

Where the random effect term *b_kl_* was omitted.

Each trait’s data was standardized to z-scores to enable comparison of effect sizes across traits. Statistical significance of fixed effects in the mixed-effects model was assessed using ANOVA, with Satterthwaite’s approximation for degree of freedom to compute p-values. For fixed-effects linear models, standard ANOVA was applied. Effect sizes (fixed-effect coefficients) and their 95% confidence intervals were extracted for both model types, providing estimates of the magnitude and direction of each factor’s influence.

### Principal Component Analysis and Random Forest regression

Following the mixed-effect modeling, a series of additional statistical analyses were conducted to investigate the relationships among the measured traits and their contributions to Grain Yield and Peak LAI. These analyses included Principal Component Analysis (PCA) and Random Forest (RF), each implemented using specific R packages and tailored to address distinct aspects of the data. PCA and RF analyses were carried out on global dataset rather than by separate treatment combinations. This decision was supported by Permutational Multivariate Analysis of Variance (PERMANOVA) using the *vegan* R package (version 2.6.10; Oksanen et al., 2022). While treatment main effects were statistically significant, interaction effect explained minimal variation in trait relationships (< 0.4% for all interaction terms), indicating that the underlying multivariate structure remained consistent across experimental conditions.

PCA was performed on traits consistently available across all experimental combinations, excluding variables with sparse data. For the variables retained for the analysis, missing values were imputed using the *mice* R package (version 3.17.0; van Buuren and Groothuis-Oudshoorn, 2011) with the predictive mean matching method (PMM), which leverages observed data patterns to estimate plausible replacements. The resulting dataset was standardized (mean = 0, standard deviation = 1) to ensure comparability across traits with differing scales. PCA was conducted using the *FactoMineR* package (version 2.11; Lê et al., 2008), employing singular value decomposition to extract principal components. Results were visualized with the *factoextra* R package (Kassambara and Mundt, 2020).

Next, RF regression was applied as a complementary approach to the multivariate analyses specifically to identify and rank the relative importance of predictor variables influencing our primary outcome measures, grain yield and peak LAI. Separate models were constructed for each of the two response variables using the *randomForest* R package (version 4.7.1.2; Liaw & Wiener, 2002), with 500 trees grown per model to ensure stability in predictions. Variable importance was assessed using two metrics: 1) the percent increase in Mean Squared Error (%IncMSE), which quantifies the increase in prediction error when a given predictor is permuted, reflecting its contribution to model accuracy; and 2) node purity, measured as the decrease in residual sum of squares (IncNodePurity), indicating a predictor’s role in partitioning the data into homogeneous subsets within the trees.

## Acknowledgements

We thank Christopher Montes and Anthony Digrado for helping with planting, Crystal Concepcion and Samuel Cheung for assisting with data collection and tissue sampling, Casey Kramer for seed protein and oil content analysis, and Sarah J. Schultz for helping develop isogenic lines.

## Author contributions

B.G.T. and E.A.A. designed the study; B.W.D. assisted in developing the population; G.B. helped in acquisition of drone imagery; C.J.B. assisted in data collection from the Spidercam^TM^ field phenotyping platform; B.G.T developed the lines, carried out the experiments, analyzed data, prepared figures and tables, and wrote the draft; E.A.A. and B.W.D edited the draft; all authors read and approved the final version of the manuscript.

---

**Figure S1.**
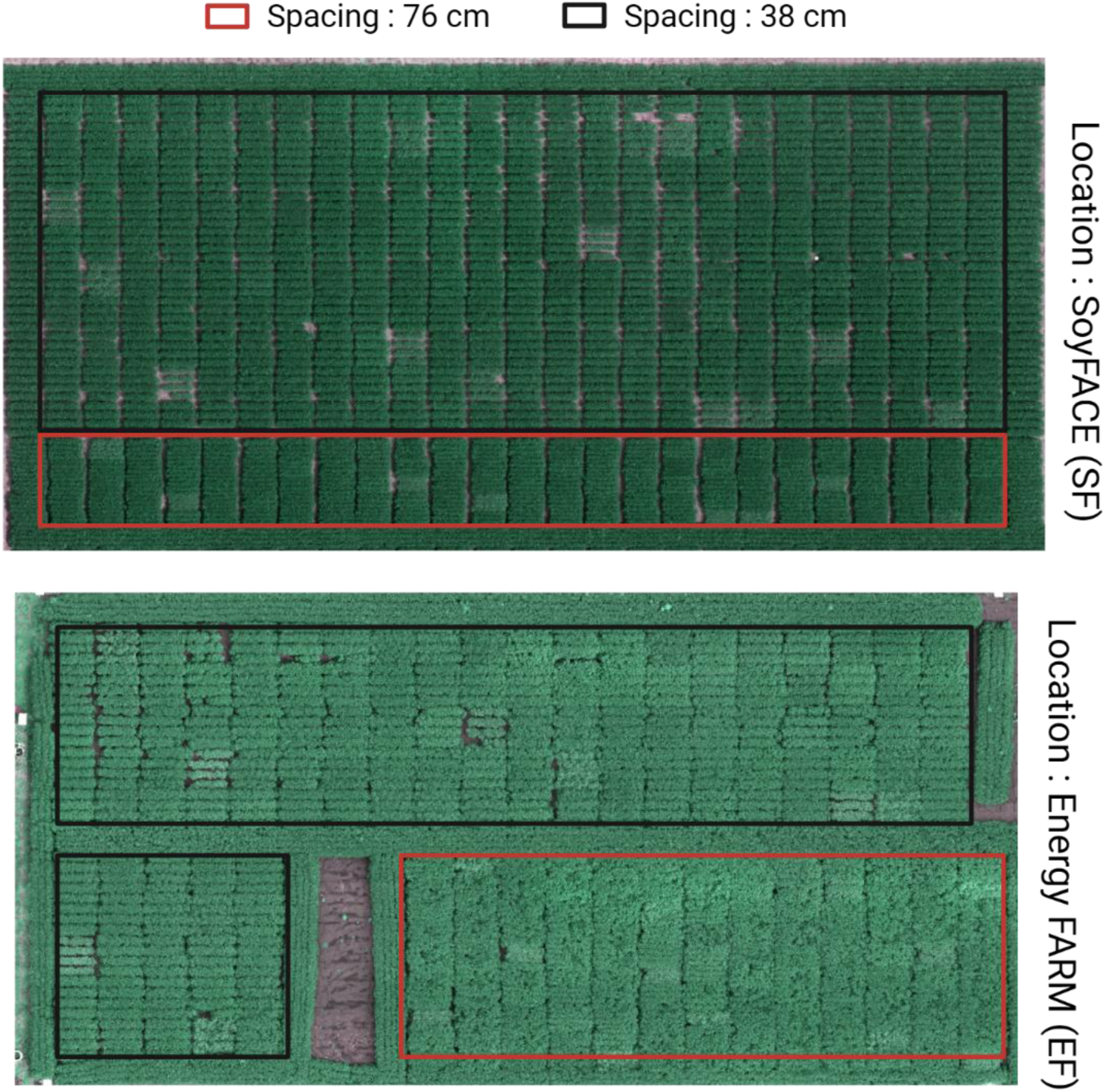
Aerial orthomosaic images of experimental plots at two field locations with contrasting row spacing treatments. **Top)** SoyFACE facility, with the 76-cm row spacing treatment demarcated by black rectangular boundaries and 38-cm row spacing by red boundaries. **Bottom)** Energy Farmfacility, located approximately 2 miles east from the SoyFACE site, with identical spacing treatments highlighted using the same color scheme. Images were captured approximately 75 days after planting when canopy closure had been achieved in most plots. Multispectral drone imagery captured during the growing season of 2024 spatial arrangement of soybean isogenic line trials.

**Figure S2.**
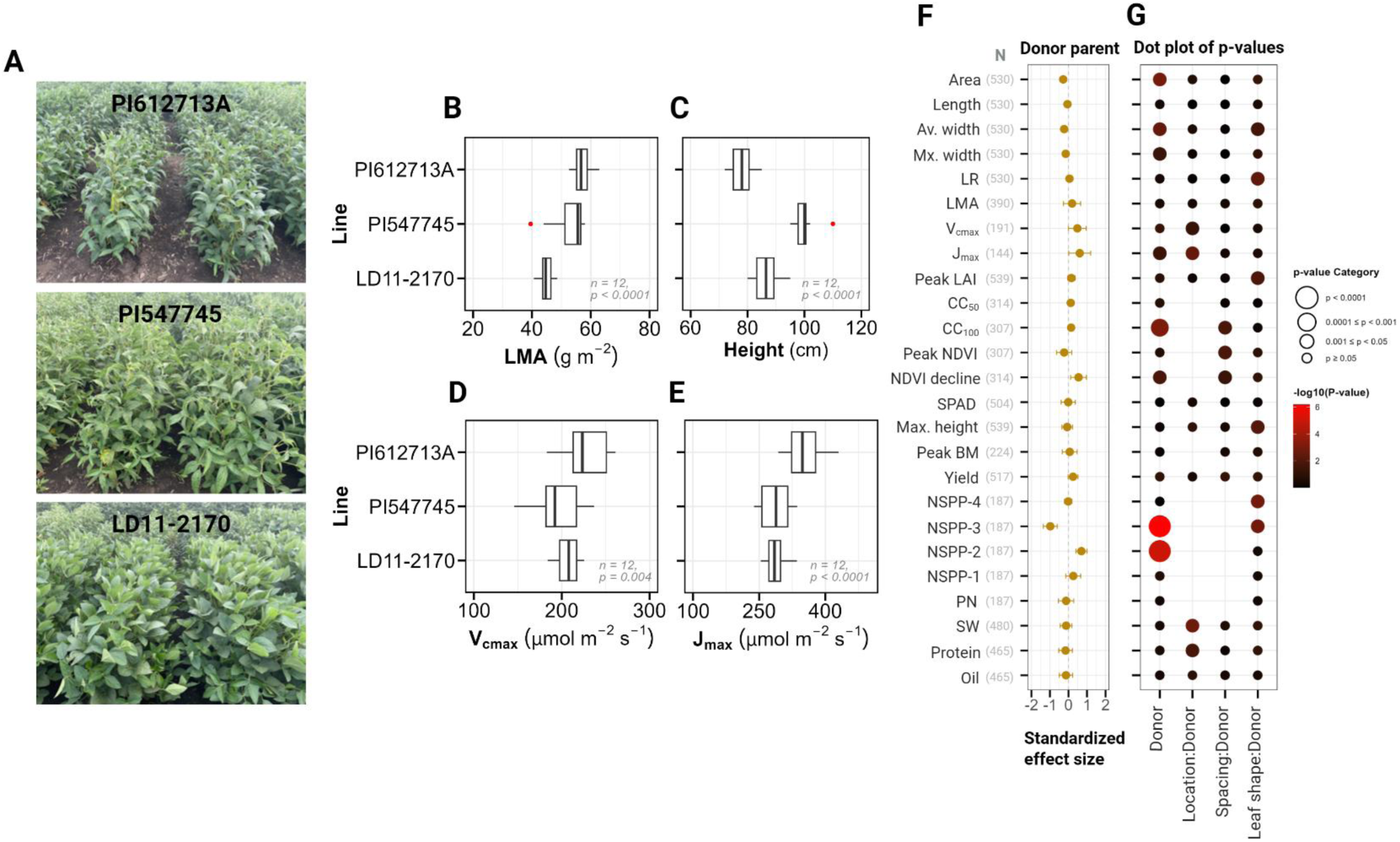
Characterization of parental lines and analysis of donor parent genetic effects on measured traits. **A)** Field images of the three parental genotypes used in developing the isogenic lines, photographed during the 2022 growing season. The narrow-leaved donor parents PI612713A (top) and PI547745 (middle) display distinctly different leaf morphology compared to the broad-leaved recurrent parent LD11-2170 (bottom). **B-E)** Box and whisker plots comparing key physiological traits across the three parental lines: Leaf Mass per Area (LMA, B), Plant height at maturity (C), maximum rate of Rubisco carboxylation (V_cmax_, D), and maximum rate of electron transport (J_max_, E). For each trait, sample size (n) and p-values from ANOVA tests are displayed within each panel. **F)** Standardized effect sizes with 95% confidence intervals (orange dots and horizontal lines) for the Donor Parent factor (PI612713A as reference) across all 25 measured traits. Sample sizes are indicated for each trait. **G)** Significance of Donor Parent and its interactions with Location, Spacing and Leaf Shape from mixed-effects modeling. The size of dots indicates significance level categories, while color intensity represents –log_10_(p-value). All effects are relative to the reference donor parent (PI612713A).

**Figure S3.**
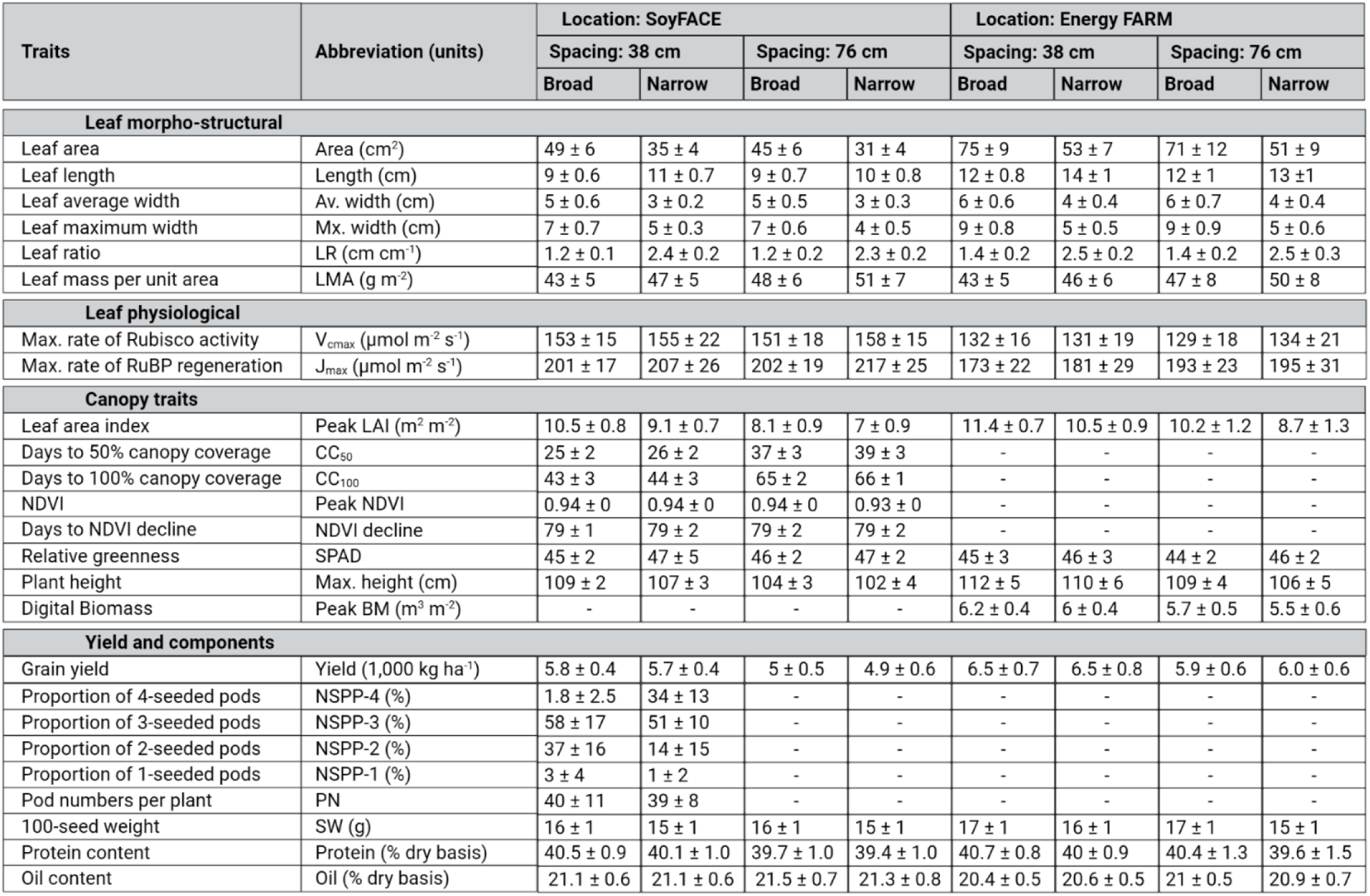
Summary statistics of morphological, physiological and agronomic traits measured across all experimental conditions. This table presents the complete set of traits assessed in this study, including standardized abbreviations, measurement units, and mean values (± SD) aggregated across both locations (SoyFACE, Energy Farm), row spacings (38-cm, 76-cm) and leaf shape (broad, narrow). These summary statistics represent data from 204 isogenic lines evaluated across 600 experimental plots in the growing season of 2024.

**Figure S4.**
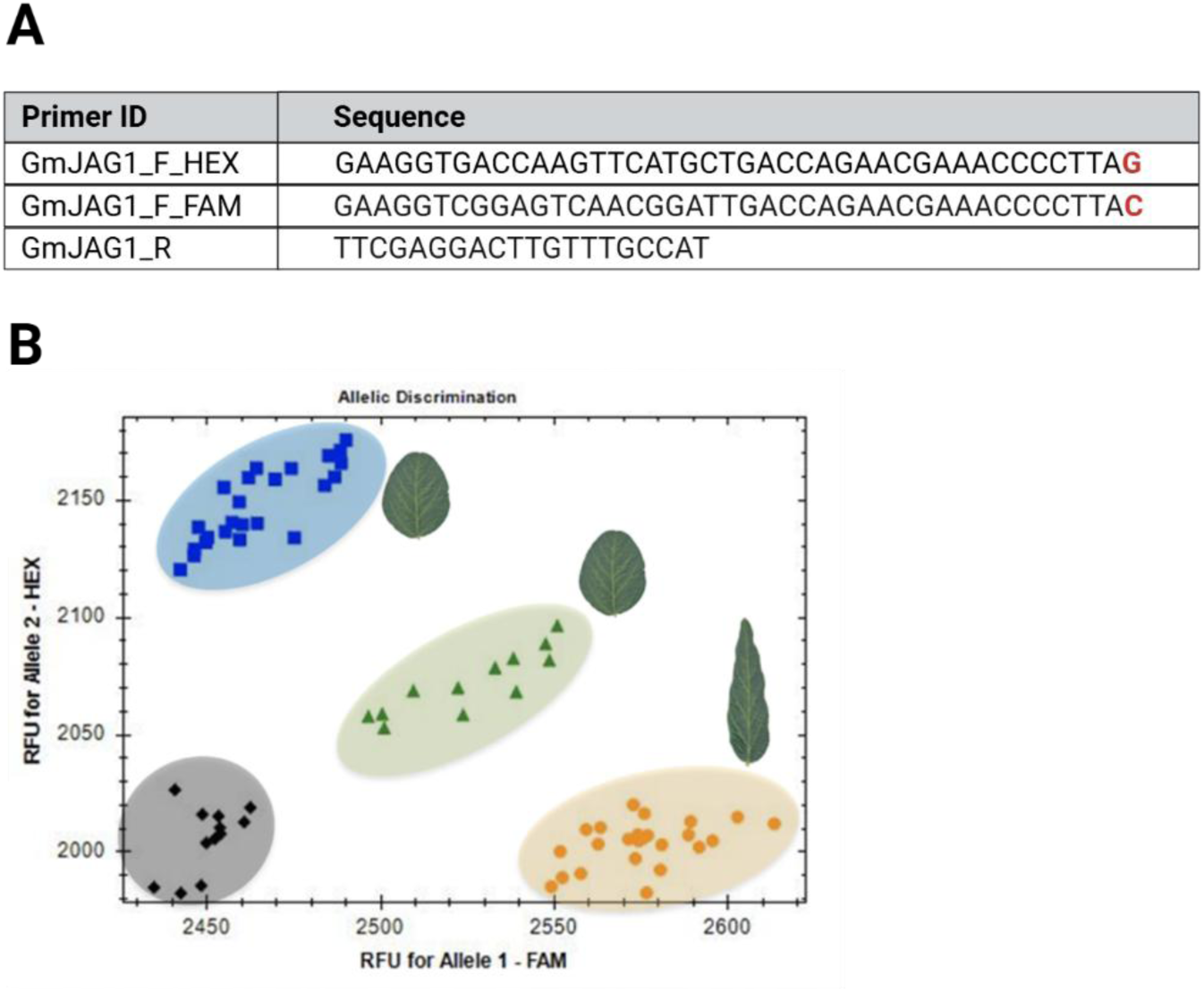
Allelic discrimination of soybean lines using KASP marker analysis. **A)** Table listing the sequences of forward (F) and reverse (R) primers used in the GmJAG1 KASP marker analysis. The allele-specific ends are highlighted in red that differentiates between the two JAG1 allele types. **B)** Scatter plot illustrating allelic discrimination of segregating soybean lines based on a Kompetitive Allele Specific PCR (KASP) marker designed for a single nucleotide polymorphism (SNP) in GmJAG1. The x-axis represents Relative Fluorescence Units (RFU) for Allele 1 (FAM) and the y-axis represents RFU for Allele 2 (HEX). Blue squares indicate homozygous dominant broad-leaved lines, orange circles represent homozygous recessive narrow-leaved lines, green triangles denote heterozygous broad-leaved lines, and black diamonds signify no-DNA-template controls. Images of broad and narrow soybean leaves are included adjacent to the corresponding blue, green and orange data points to visually distinguish leaf morphologies.

**Figure S5.**
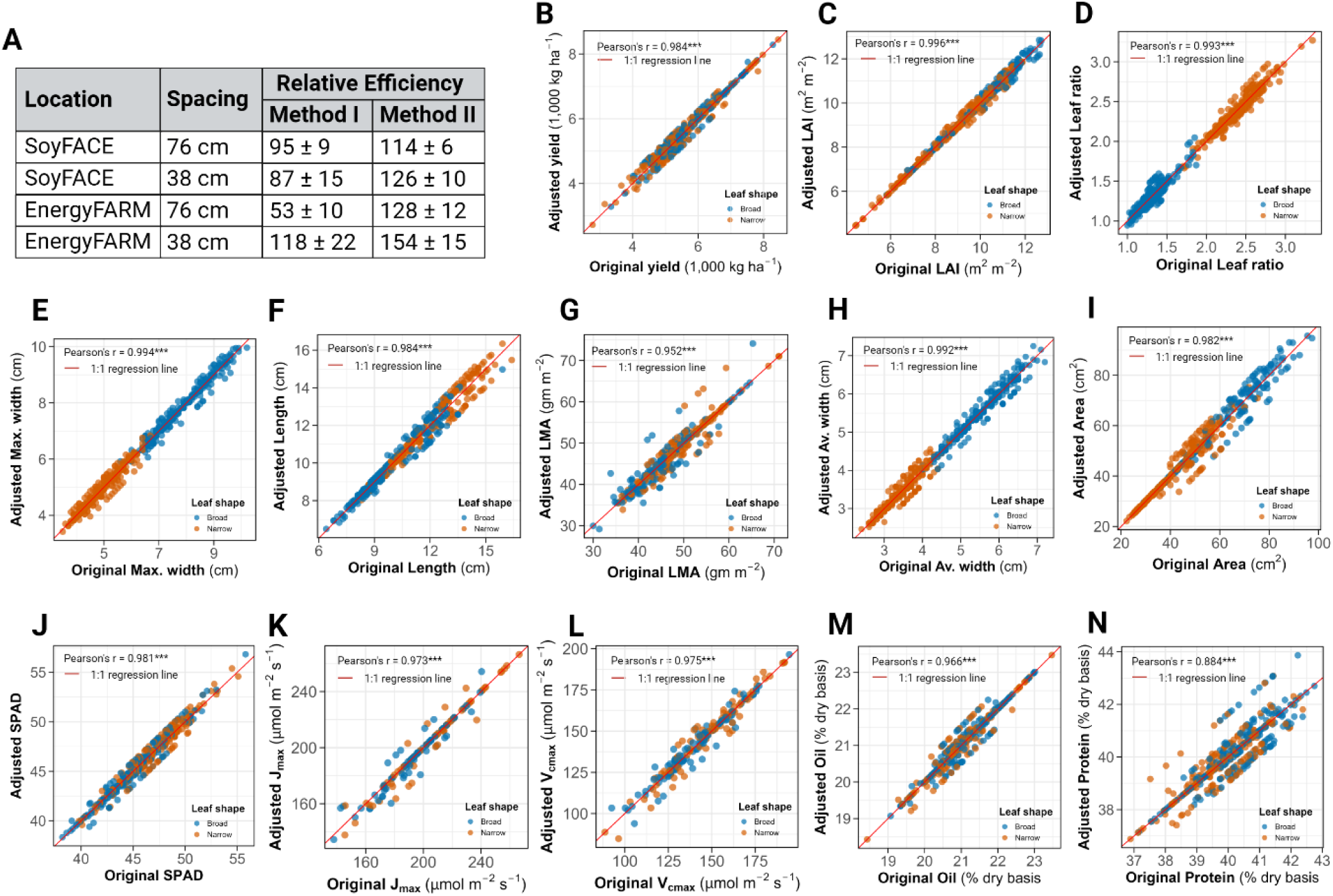
Relative Efficiency (RE) analysis of Type II Modified Augmented Design. **A)** Relative efficiency comparison between Method I and Method II across two locations and two spacings. Values represent the mean RE ± standard deviation calculated across all evaluated traits. **B-N)** Correlation between original and adjusted values for the traits which required adjustments. Data points are categorized by leaf shape, Broad (blue) and Narrow (orange). The red line represents the line 1:1 regression line. Pearson’s correlation coefficients (r) are indicated with significance levels: * p < 0.05, ** p < 0.01, *** p < 0.001.

**Figure S6.**
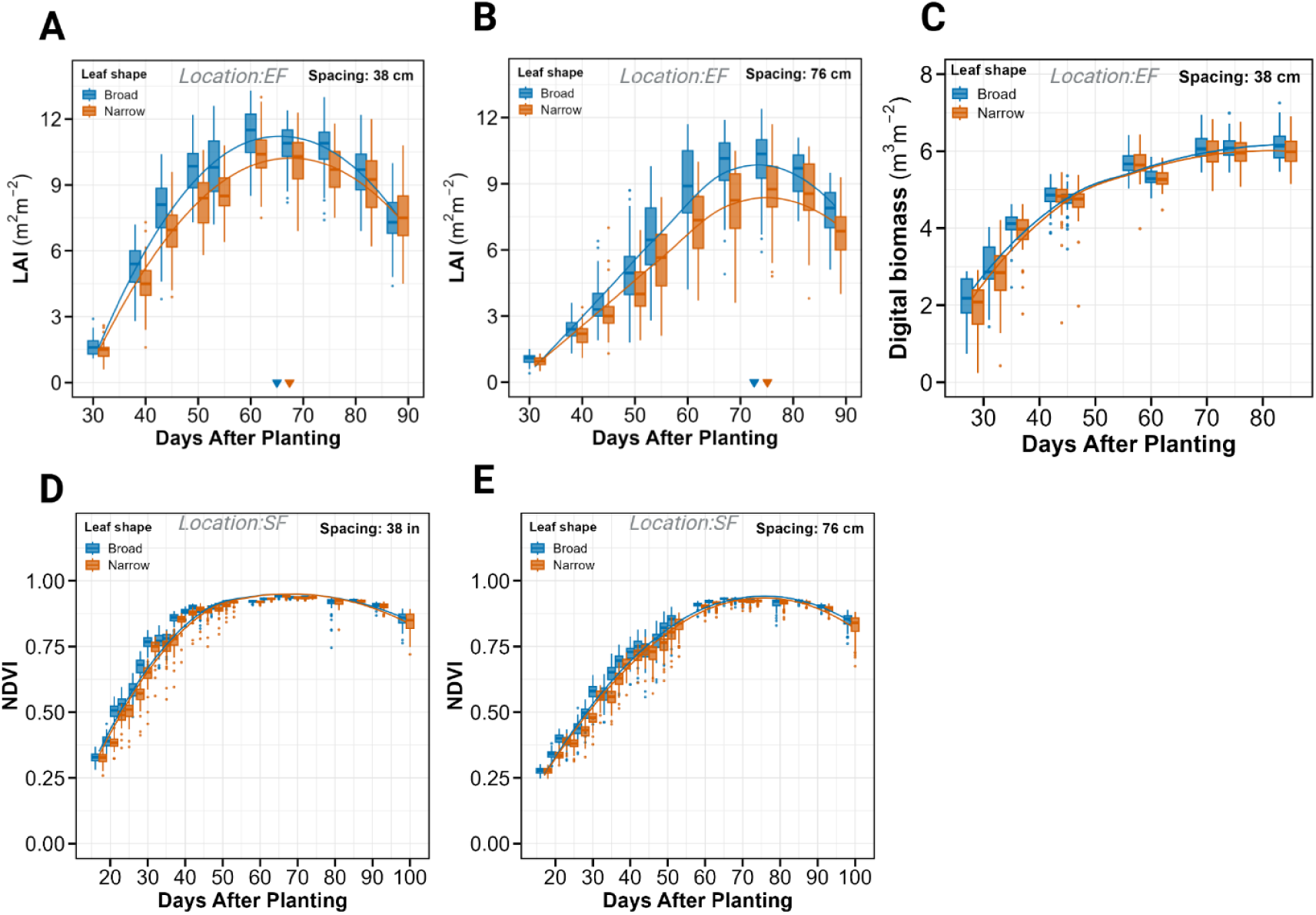
Temporal dynamics of canopy development traits. **A-B)** Leaf Area Index (LAI) time course at Energy Farm at 38-cm and 76-cm row spacings. Each time-point displays box and whisker plots for broad- (blue) and narrow-leaved (orange) lines. Box plots display the median, interquartile range (IQR), and whiskers represent 1.5 times IQR with outliers shown as individual points. LOWESS curves illustrate the temporal trends for each leaf morphology. Triangles on the x-axis indicate model-predicted peak LAI values. **C)** Digital biomass progression over time at the Energy Farm at 38-cm row spacing. Data visualization follows the same format as panels A-B, with box and whisker plots for broad and narrow-leaved lines at each time point, fitted with LOWESS curves. **D-E)** Normalized Difference in Vegetation Index (NDVI) time course at SoyFACE in 38-cm and 76-cm row spacings. Data presentation follows the same format as panel C, showing the temporal dynamics of NDVI for both leaf morphologies.

**Figure S7.**
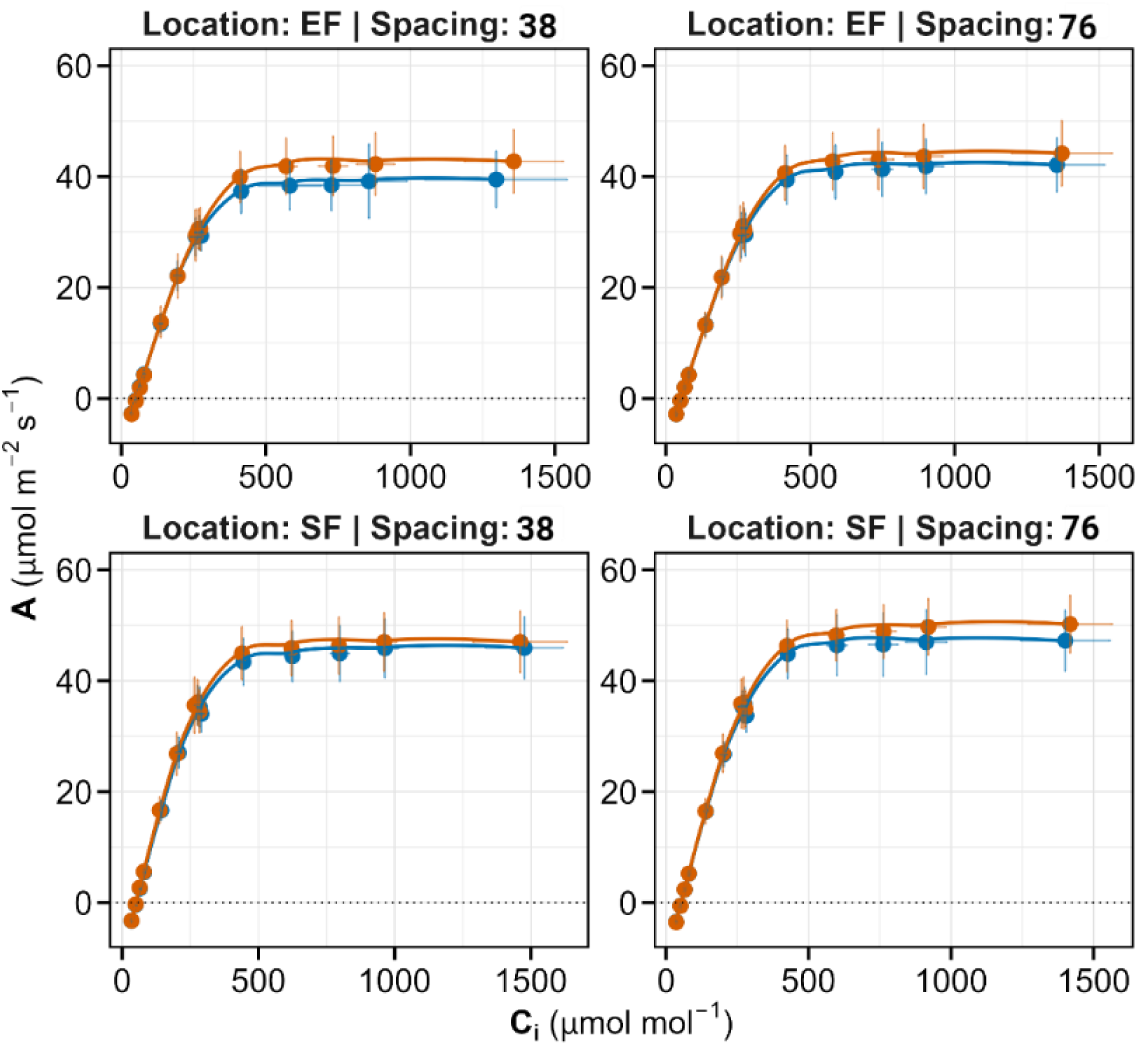
A-C_i_ response curves at different locations and row spacings. The four panels show the relationship between intercellular CO_2_ concentration (C_i_) and photosynthetic rate (A) for broad- and narrow-leaved lines. Each data point represents the mean value with standard deviation bars in both axes. Fitted curves illustrate the photosynthetic response patterns for each leaf morphology.

**Figure S8.**
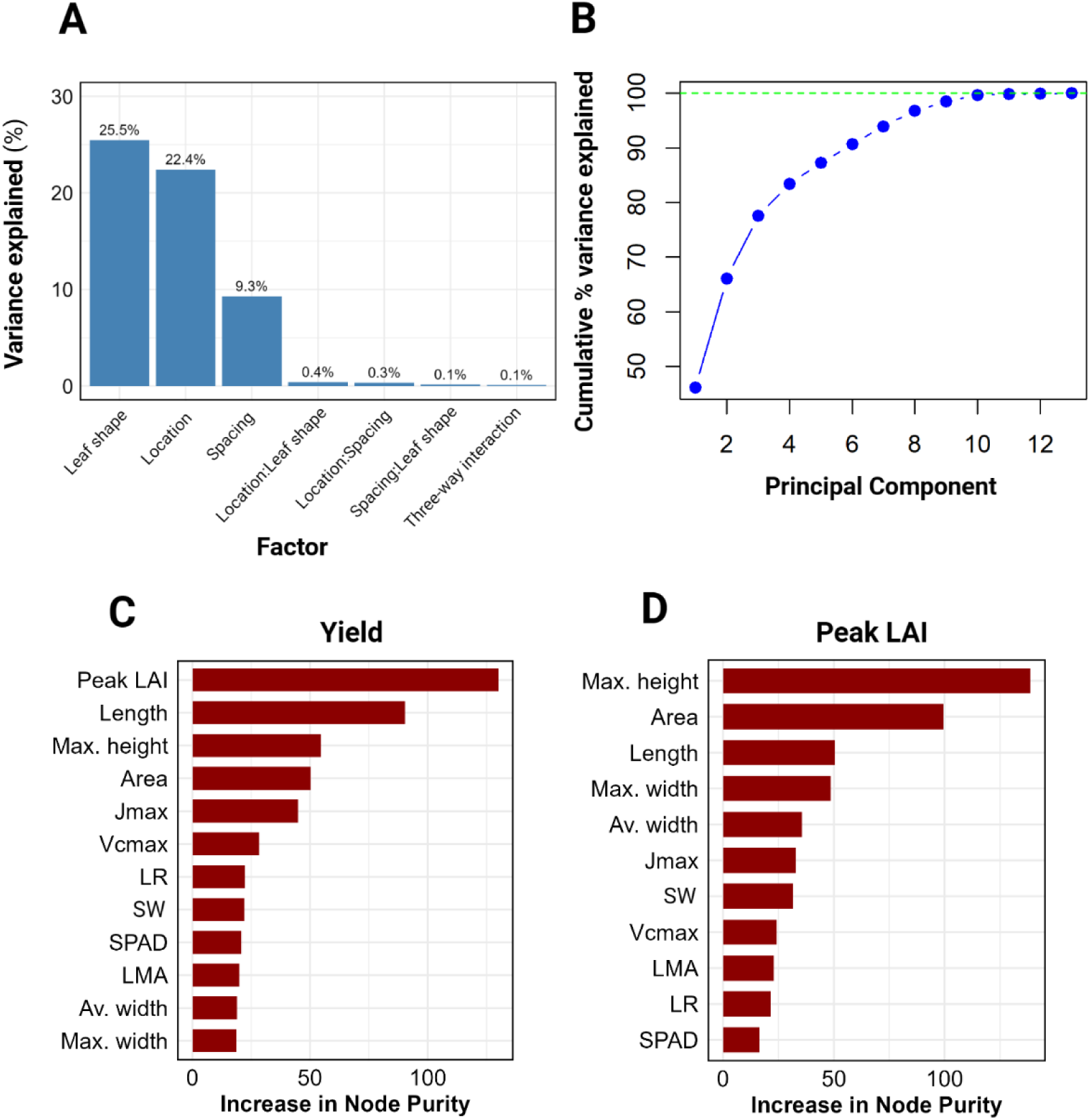
Multivariate analysis of experimental factors and traits. **A)** PERMANOVA results showing the variance explained by the three experimental factors: Location, Spacing and Leaf shape and their interactions. Blue bars represent the percentage of variance explained by each factor, with values displayed at the top of each bar. **B)** Cumulative variance explained by principal components, illustrating the contribution of each additional principal component to the total explained variance in the dataset. **C-D)** Random Forest Variable Importance analysis for Yield and Peak LAI. Red bars represent the increase in node purity for each predictor variable, indicating their relative importance in predicting the response variables.

**Figure S9.**
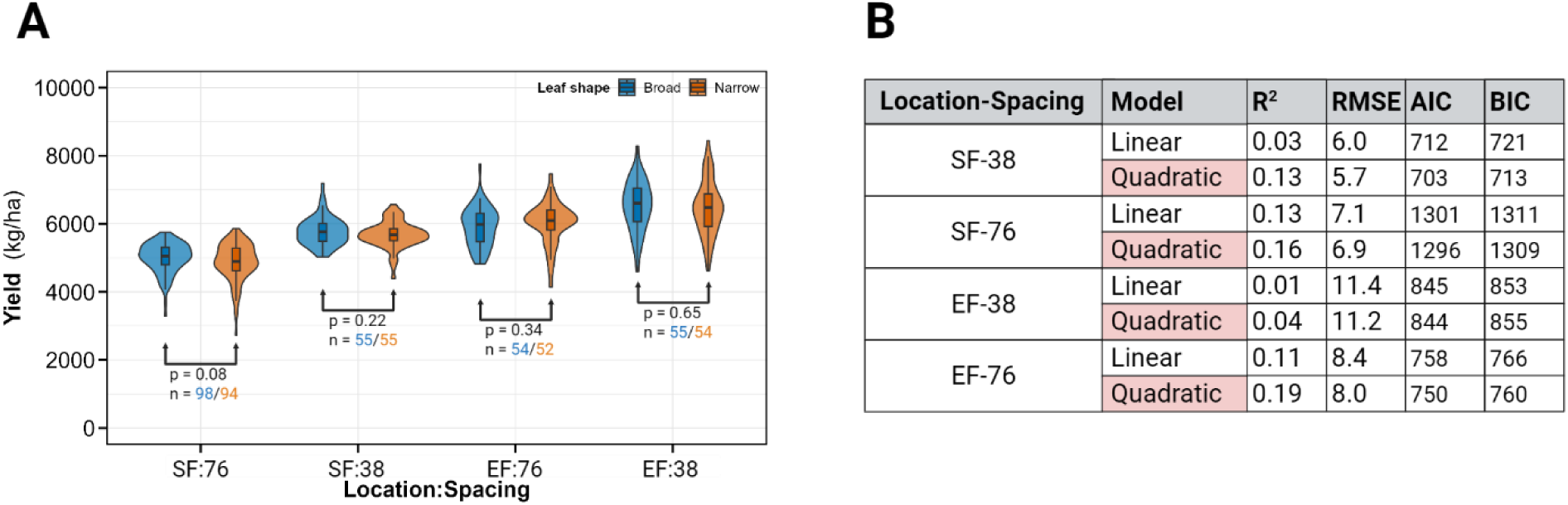
Yield comparison between leaf morphologies and model assessment of Peak LAI vs. Yield relationship. **A)** Violin plots comparing yield performance between broad-leaved and narrow-leaved soybean isogenic lines across four locations and spacing combinations. The embedded box plots display the median, interquartile range (IQR), and whiskers represent 1.5 times IQR. Statistical significance was assessed using t-tests, with corresponding p-values and sample sizes (n) indicated for each comparison. **B)** Table summarizing the linear and quadratic regression model fits for the relationship between Peak LAI and Yield across the same four location and spacing combinations. For each model, R-squared (R^2^), Root Mean Squared Error (RMSE), Akaike Information Criterion (AIC) and Bayesian Information Criterion (BIC) are reported to evaluate model performance, with the quadratic model providing the estimated Peak LAI at maximum yield.

**Figure S10.**
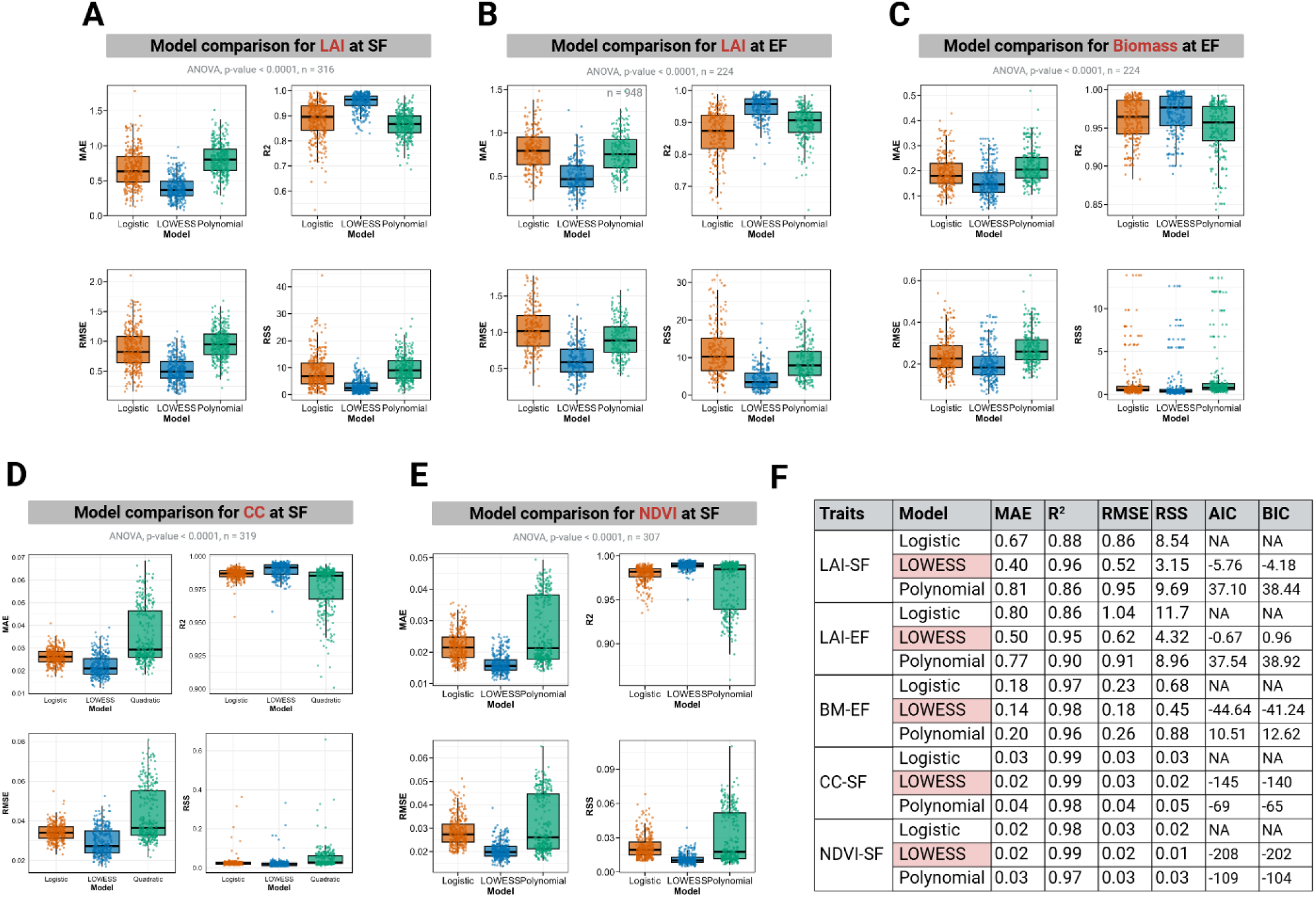
Model performance comparison across multiple canopy traits. Box and whisker plots comparing model fit metrics (MAE, R^2^, RMSE and RSS) for three curve-fitting approaches: Logistic (orange), LOWESS (blue) and Polynomial (green). Individual data-points represent experimental units. ANOVA p-values and sample sizes (n) are provided for each comparison. **A-B)** Model performance metrics for Leaf Area Index at SoyFACE and Energy Farm, respectively. **C)** Model performance metrics for Digital Biomass at Energy Farm. **D)** Model performance metrics for Canopy Coverage at SoyFACE. **E)** Model performance metrics for NDVI at SoyFACE. **F)** Summary table of mean values for traditional fit metrics (MAE, R^2^, RMSE, RSS) and information criteria (AIC, BIC) across all assessed traits and models. Values represent means across all experimental units, combining both row spacings. Lower values of MAE, RMSE, RSS, AIC and BIC indicate better model performance, while higher R^2^ values indicate better model fit.

## References

Ainsworth EA, Long SP. What have we learned from 15 years of free-air CO₂ enrichment (FACE)? A meta-analytic review of the responses of photosynthesis, canopy properties and plant production to rising CO₂. New Phytologist. 2005:165:351– 372.

Andrade FH, Calvino P, Cirilo A, Barbieri P. Yield responses to narrow rows depend on increased radiation interception. Agronomy Journal. 2002:94:975–980.

Andrade JF, Edreira JIR, Mourtzinis S, Conley SP, Ciampitti IA, Dunphy JE, Gaska JM, Glewen K, Holshouser DL, Kandel HJ, et al. Assessing the influence of row spacing on soybean yield using experimental and producer survey data. Field Crops Research. 2019:230:98–106.

Bailey-Serres J, Parker JE, Ainsworth EA, Oldroyd GED, Schroeder JI. Genetic strategies for improving crop yields. Nature. 2019:575:109–118.

Bates D, Maechler M, Bolker B, Walker S. Fitting linear mixed-effects models using lme4. Journal of Statistical Software. 2015:67:1–48.

Bernachhi CJ, Singsaas EL, Pimentel C, Portis JR AR, Long SP. Improved temperature response functions for models of Rubisco-limited photosynthesis. Plant, Cell & Environment. 2001:24:253–259.

Bianchi JS, Quijano A, Gosparini CO, Morandi EN. Changes in leaflet shape and seeds per pod modify crop growth parameters, canopy light environment, and yield components in soybean. The Crop Journals. 2020:8:351–364.

Bloom AJ, Chapin FS III, Mooney HA. Resource limitation in plants-An economic analogy. Annual Review of Ecology and Systematics. 1985:16:363–392.

Board JE, Harville BG, Saxton AM. Narrow-row seed-yield enhancement in determinate soybean. Agronomy Journal. 1990:82:64–68.

Board JE, Harville BG. Explanations for greater light interception in narrow-vs. wide-row soybean. Crop Science. 1992:32:198–202.

Busch FA, Ainsworth EA, Amtmann A, Cavanagh AP, Driever SM, Ferguson JN, Kromdijk J, Lawson T, Leakey ADB, Matthews JSA, et al. A guide to photosynthetic gas exchange measurements: Fundamental principles, best practice and potential pitfalls. Plant, Cell & Environment. 2024:47:3344–3364.

Cai G, Brock A. The uniform soybean tests, northern region 2023. USDA-ARS. 2023. https://www.ars.usda.gov/ARSUSERFILES/50200500/UST/2023.PDF

Cai Z, Xian P, Cheng Y, Ma Q, Lian T, Nian H, Ge L. CRISPR/Cas9-mediated gene editing of GmJAGGED1 increased yield in the low-latitude soybean variety Huachun 6. Plant Biotechnology Journal. 2021:19:1898–1900.

Chen Y, Nelson RL. Evaluation and classification of leaflet shape and size in wild soybean. Crop Science. 44:671–677.

Clark CB, Ma J. The genetic basis of shoot architecture in soybean. Molecular Breeding. 2023:43:55.

Croce R, Carmo-Silva E, Cho YB, Ermakova M, Harbinson J, Lawson T, McCromick AJ, Niyogi KK, Ort DR, Patel-Tupper D, et al. Perspectives on improving photosynthesis to increase crop yield. The Plant Cell. 2024:36:3944–3973.

Digrado A, Ainsworth EA. Modifying canopy architecture to optimize photosynthesis in crops. In: Understanding and improving crop photosynthesis, 1st ed. Burleigh Dodds Science Publishing. 2023:1–42.

Digrado A, Mitchell NG, Montes CM, Dirvanskyte P, Ainsworth EA. Assessing diversity in canopy architecture, photosynthesis, and water-use efficiency in a cowpea magic population. Food and Energy Security. 2020:9:e236.

Dinkins RD, Keim KR, Farno L, Edwards LH. Expression of the narrow leaflet gene for yield and agronomic traits in soybean. The Journal of Heredity. 2002:93:346– 351.

Evans JR. Photosynthesis and nitrogen relationships in leaves of C₃ plants. Oecologia. 1989:78:9–19.

FAO. Agricultural production statistics 2000-2022. FAOSTAT Analytical Briefs, No. 79. Rome. 2023. 10.4060/cc9205en

Fehr WR, Caviness CE, Burmood DT, Pennington JS. Stage of development descriptions for soybeans, *Glycine max* (L.) Merrill. Crop Science. 1971:11:929–931.

Flood PJ, Harbinson J, Aarts MGM. Natural genetic variation in plant photosynthesis. Trends in Plant Science. 2011:16:327–335.

Goulart HMD, van der Wiel K, Folberth C, Boere E, van den Hurk B. Increase of simultaneous soybean failures due to climate change. Earth’s Future. 2023:11:e2022EF003106.

Holland BL, Monk NAM, Clayton RH, Osborne CP. A theoretical analysis of how plant growth is limited by carbon allocation strategies and respiration. In Silico Plants. 2019:1:diz004.

Jeong N, Suh SJ, Kim M-H, Lee S, Loon J-K, Kim HS, Jeong S-C. Ln is a key regulator of leaflet shape and number of seeds per pod in soybean. The Plant Cell. 2012:24:4807–4818.

John GP, Garnica-Diaz CJ. Embracing the complexity of leaf shape: a commentary on ‘Anatomical determinants of gas exchange and hydraulics vary with leaf shape in soybean. Annals of Botany. 2023:131:i–iii.

Kassambara A, Mundt F. Factoextra: Extract and Visualize Results of Multivariate Data Analyses. R package version 1.0.7. 2020. https://CRAN.R-project.org/package=factoextra.

Krause MD, Dias KOG, Singh AK, Beavis WD. Using soybean historical field trial data to study genotype by environment variation and identify mega-environments with the integration of genetic and non-genetic factors. Agronomy Journal. 2025:117:e70023.

Kuznetsova A, Brockhoff PB, Christensen RHB. lmerTest Package: Tests in Linear Mixed Effects Models. Journal of Statistical Software. 2017:82:1–26.

Lê S, Josse J, Husson F. FactoMineR: An R package for multivariate analysis. Journal of Statistical Software. 2008:25:1–18.

Length R. Emmeans: estimated marginal means, aka least-squares means. 1.10.7. 2025. https://CRAN.R-project.org/package=emmeans

Li W, Wang L, Xue H, Zhang M, Song H, Qin M, Dong Q. Molecular and genetic basis of plant architecture in soybean. Frontiers in Plant Science. 2024:15:1477616.

Liaw A, Wiener M. Classification and regression by randomForest. R News. 2002:2:18– 22.

Lin C-S, Poushinsky G. A modified augmented design (type 2) for rectangular plots. Canadian Journal of Plant Sciences. 1985:65:743–749.

Liu S, Baret F, Abichou M, Manceau L, Andrieu B, Weiss M, Martre P. Importance of the description of light interception in crop growth models. Plant Physiology. 2021:186:977–997.

Liu X, Jin J, Wang G, Herbert SJ. Soybean yield physiology and development of high-yielding practices in Northeast China. Field Crops Research. 2008:105:157–171.

Lochocki E, Salesse-Smith CE, McGrath JM. PhotoGEA: An R package for closer fitting of photosynthetic gas exchange data with non-gaussian confidence interval estimation. Plant, Cell & Environment. 2025:1–16.

Long S, Amy M-C, Zhu X-G. Meeting the global food demand of the future by engineering crop photosynthesis and yield potential. Cell. 2015:161:56–66.

Long SP, Bernacchi CJ. Gas exchange measurements, what can they tell us about the underlying limitations to photosynthesis? Procedures and sources of error. Journal of Experimental Botany. 2003:54:2392–2401.

Lu F, Hongyan W, Xiaowei M, Hongbo P, Jianrong S. Modeling the current land suitability and future dynamics of global soybean cultivation under climate change scenarios. Field Crops Research. 2021:263:108069.

Mandl FA, Buss GR. Comparison of narrow and broad leaflet isolines in soybean. Crop Science. 1981:21:25–27.

Mantilla-Perez MB, Fernandez MGS. Differential manipulation of leaf angle throughout the canopy: current status and prospects. Journal of Experimental Botany. 2017:68:5699–5717.

Messina M. Perspective: Soybeans Can Help Address the Caloric and Protein Needs of a Growing Global Population. Frontiers in Nutrition. 2022:9:909464.

Miflin B. Crop improvement in the 21st century. Journal of Experimental Botany. 2000:51:1–8.

Moreau D, Allard V, Gaju O, Le Gouis J, Foulkes MJ, Martre P. Acclimation of leaf nitrogen to vertical light gradient at anthesis in wheat is a whole-plant process that scales with the size of the canopy. Plant Physiology. 2012:160:1479–1490.

Niinemets U. A review of light interception in plant stands from leaf to canopy in different plant functional types and in species with varying shade tolerance. Ecological Research. 2010:25:693–714.

Nobel PS, Forseth IN, Long SP. Canopy structure and light interception. In: Hall DO, Scurlock JMO, Bolhar-Nordenkampf HR, Leegood RC, Long SP, editors. Photosynthesis and Production in a Changing Environment: a field and laboratory manual. Chapman and Hall, London. 1993:79–90.

Oksanen J, Simpson GL, Blanchet FG, Kindt R, Legendre P, Minchin PR, O’Hara RB, Solymos P, Stevens MHH, Szoecs E, et al. vegan: community ecology package. 2.6.10. 2022. https://CRAN.R-project.org/package=vegan

Onoda Y, Wright IJ, Evans JR, Hikosaka K, Kitajima K, Niinemets U, Poorter H, Tosens T, Westoby M. Physiological and structural tradeoffs underlying the leaf economic spectrum. New Phytologist. 2017:214:1447–1463.

Parker GG. Tamm review: Leaf Area Index (LAI) is both a determinant and a consequence of important processes in vegetation canopies. Forest Ecology and Management. 2020:477:118496.

Poorter H, Evans JR. Photosynthetic nitrogen-use efficiency of species that differ inherently in specific leaf area. Oecologia. 1998:116:26–37.

Poorter H, Niinemets U, Poorter L, Wright IJ, Villar R. Causes and consequences of variation in leaf mass per area (LMA): a meta-analysis. New Phytologist. 2009:182:565–588.

Purcell LC, Ball RA, Reaper JD III, Vories ED. Radiation use efficiency and biomass production in soybean at different population densities. Crop Science. 2002:42:172–177.

Qiang B, Zhou W, Zhong X, Fu C, Cao L, Zhang Y, Jin X. Effect of nitrogen application levels on photosynthetic nitrogen distribution and use efficiency in soybean seedling leaves. Journal of Plant Physiology. 2023:287:154051.

R Core Team. R: a language and environment for statistical computing. R foundation for statistical computing, Vienna, Austria. 2024. https://www.R-project.org/.

Richards RA. Selectable traits to increase crop photosynthesis and yield of grain crops. Journal of Experimental Botany. 2000:51:447–458.

Roth L, Barendregt C, Betrix C-A, Hund A, Walter A. High-throughput field phenotyping of soybean: spotting an ideotype. Remote Sensing of Environment. 2022:269:112797.

Rusu RB, Cousins S. 3D is here: Point Cloud Library (PCL). In: IEEE International Conference on Robotics and Automation (ICRA). Shanghai, China. 2011.

Sakamoto T, Matsuoka M. Generating high-yielding varieties by genetic manipulation of plant architecture. Current Opinion in Biotechnology. 2004:15:144–147.

Sarlikioti V, de Visser PHB, Marcelis LFM. Exploring the spatial distribution of light interception and photosynthesis of canopies by means of a functional-structural plant model. Annals of Botany. 2011:107:875–883.

Schiessl K, Muino JM, Sablowski R. Arabidopsis JAGGED links floral organ patterning to tissue growth by repressing Kip-related cell cycle inhibitors. Proceedings of the National Academy of Sciences. 2014:111:2830–2835.

Setiyono TD, Weiss A, Specht JE, Cassman KG, Dobermann A. Leaf area index simulation in soybean grown under near-optimal conditions. Field Crops Research. 2008:108:82–92.

Singh M, Thapa R, Singh N, Mirsky SB, Acharya BS, Jhala AJ. Does narrow row spacing suppress weeds and increase yields in corn and soybean? A meta-analysis. Weed Science. 2023:71:520–535.

Slattery RA, Ort DR. Perspectives on improving light distribution and light use efficiency in crop canopies. Plant Physiology. 2021:185:34–48.

Soares JC, Zimmermann L, dos Santos NZ, Muller O, Pintado M, Vasconcelos MW. Genotypic variation in the response of soybean to elevated CO₂. Plant-Environment Interaction. 2021:2:263–276.

Sreekanta S, Haaning A, Dobbels A, O’Neil R, Hofstad A, Virdi K, Katagiri F, Stupar RM, Muehlbauer GJ, Lorenz AJ. Variation in shoot architecture traits and their relationship to canopy coverage and light interception in soybean (*Glycine max*). BMC Plant Biology. 2024:24:194.

Srinivasan V, Kumar P, Long SP. Decreasing, not increasing, leaf area will raise crop yields under global atmospheric change. Global Change Biology. 2017:23:1626–1635.

Stupar RM, Locke AM, Allen DK, Stacey MG, Ma J, Weiss J, Nelson RT, Hudson ME, Joshi T, Li Z, et al. Soybean genomic research community strategic plan: A vision for 2024-2028. The Plant Genome. 2024:17:e20516.

Tagliapietra EL, Streck NA, da Rocha TSM, Richter GL, da Silva MR, Cera JC, Guedes JVC, Zanon AJ. Optimum leaf area index to reach soybean yield potential in subtropical environment. Crop Ecology and Physiology. 2018:110:932–938.

Tamang BG, Zhang Y, Zambrano MA, Ainsworth EA. Anatomical determinants of gas exchange and hydraulics vary with leaf shape in soybean. Annals of Botany. 2023:131:909–920.

van Buuren S, Groothuis-Oudshoorn K. Mice: multivariate imputation by chained equations in R. Journal of Statistical Software. 2011:45:1–67.

Villar R, Olmo M, Atienza P, Garzon AJ, Wright IJ, Poorter H, Hierro LA. Applying the economic concept of profitability of leaves. Scientific Reports. 2021:11:49.

Walker AP, Beckerman AP, Gu L, Kattge J, Cernusak LA, Domingues TF, Scales JC, Wohlfahrt G, Wullschleger SD, Woodward FI. The relationship of leaf photosynthetic traits -- Vcmax and Jmax -- to nitrogen, leaf phosphorus, and specific leaf area: a meta-analysis and modeling study. Ecology and Evolution. 2014:4:3218–3235.

Webster RW, Rother MG, Mueller BD, Mueller DS, Chilvers MI, Willbur JF, Mourtzinis S, Conley SP, Smith DL. Integration of row spacing, seeding rates, and fungicide applications for control of sclerotinia stem rot in *Glycine max*. Plant Disease. 2022: 106:1183–1191.

Wei B, Ma X, Guan H, Yu M, Yang C, He H, Wang F, Shen P. Dynamic simulation of leaf area index for the soybean canopy based on 3D reconstruction. Ecological Informatics. 2023:75:102070.

Wells R, Burton JW, Kilen TC. Soybean growth and light interception: response to differing leaf and stem morphology. Crop Science. 1993:33:520–524.

Wickham H, Francois R, Henry L, Muller K, Vaughan D. dplyr: a grammar of data manipulation. R package version 1.1.4. 2023. https://CRAN.R-project.org/package=dplyr

Wickham H, Vaughan D, Girlich M. tidyr: tidy messy data. R package version 1.3.1. 2023. https://CRAN.R-project.org/package=tidy.

Wickham H. ggplot2: Elegant Graphics for Data Analysis. Springer-Verlag New York. 2016. ISBN 978-3-319-24277-4. https://ggplot2.tidyverse.org/

Wright IJ, Reich PB, Westoby M, Ackerly DD, Baruch Z, Bongers F, Cavender-Bares J, Chapin T, Cornelissen JHC, Diemer M, et al. The worldwide leaf economic spectrum. Nature. 2004:428:821–827.

Yang X, Li R, Jablonski A, Stovall A, Kim J, Yi K, Ma Y, Beverly D, Phillips R, Novick K, et al. Leaf angle as a leaf and canopy trait: Rejuvenating its role in ecology with new technology. Ecology Letters. 2023:26:1005–1020.

